# Stress Granules Buffers Inflammation by Restricting dsRNA-led Mitochondrial Fragmentation

**DOI:** 10.64898/2026.03.23.713569

**Authors:** Prerna Narwal, Shovon Swarnakar, K Shreya, Narjis Fatima, Prarthana Ghosh Dastidar, Akshay Lonare, Jitendra Singh, Abhinav Banerjee, Mahipal Ganji, Jomon Joseph, Jaydeep K. Basu, Shovamayee Maharana

## Abstract

Stress granules (SGs) are dynamic RNA–RBP condensates that form during stress and inflammation, yet how they modulate inflammatory signalling remains unclear. We uncover a rapid, protective SG-mediated mechanism that preserves mitochondrial integrity. During stress and translation inhibition, mitochondrial fragmentation releases double-stranded RNA (dsRNA), which activates PKR and its downstream effector DRP1, generating a self-amplifying loop of mitochondrial fragmentation and inflammation. We find that released dsRNA nucleates nanoSGs within minutes at ER–mitochondria contact sites—the very sites of mitochondrial division. These nanoSGs grow into macroSGs, effectively sequestering PKR-activating dsRNA from the cytosol. By depleting dsRNA, SGs suppress PKR–DRP1–driven positive-feedback inflammation and maintain mitochondrial integrity and function. Our findings reveal SGs as key guardians of mitochondrial homeostasis and position condensate biology at the centre of chronic mitochondrial-driven inflammation relevant to autoimmunity, ageing, and neurodegenerative disease.

---

Cells experience diverse stresses arising from fluctuations in nutrients, energy, temperature, and pathogenic infections. To cope, cells rapidly reprogram gene expression, including global repression of housekeeping genes and induction of stress-responsive genes at transcriptional and translational levels. Translation shutdown promotes formation of cytoplasmic RNA–RBP condensates called stress granules (SGs), which contain ribosomes (*1*), translation regulators, scaffold RBPs (*2*), and multiple RNA species (*3*), (*4*). SGs are implicated in RNA protection and buffering inflammation. They also associate with membrane-bound organelles such as lysosomes for maintaining its integrity (*5*), the ER where they cluster IRE1α to activate pro-survival signaling (*6*), and the Golgi through G3BPs and RNA (*7*). SGs additionally interact with mitochondria during starvation, influencing mitochondrial metabolism and ROS production (*8*). These findings highlight emerging roles of SG–organelle interactions, though their functions under stress and physiological conditions remain unclear.

Mitochondria are central to stress, chronic inflammation, ageing, and neurodegeneration. Oxidative (*9*), heat (*10*), and infection-related stresses (bacterial (*11*); viral (*12*)) promote mitochondrial fragmentation, as does ageing through altered fission–fusion dynamics (*13*). Stress and ageing also increase mitochondrial permeability via VDAC (*14*), (*15*) and BAX/BAK macropores (*16*). These structural changes enhance release of ROS, pro-apoptotic factors (*17*), oxidised DNA (*15*), dsDNA (*14*), and dsRNA (*16*) all of which drive inflammation. SGs buffer inflammation by sequestering dsRNA (*18*) and dsRNA-sensing proteins, including RIG-I, MAVS, OAS, and PKR (*19*). The coexistence of dsRNA-sequestering SGs and mitochondrial stress responses suggests a functional link, further supported by detection of mitochondrial rRNAs (mt-rRNA), mt-tRNAs, and mt-mRNAs within SGs (*3, 4, 20, 21*), though no mechanistic connection has been established.

To define this connection, we investigated mitochondrial transcripts in SGs. We found that stress and translation inhibition drive mitochondrial dsRNA enrichment in SGs. Live-cell imaging revealed that mitochondrial transcription accelerates SG nucleation. Translation inhibition alone elevated mitochondrial dsRNA and activated PKR, initiating a positive feedback loop that promoted mitochondrial fragmentation and dsRNA release through DRP1 activation at Endoplasmic Reticulum Mitochondrial Contact Sites (ERMCSs). The released dsRNA triggered the rapid formation of nanoSGs within minutes, which subsequently grew into macroSGs and were sufficient to prevent mitochondrial fragmentation and buffer inflammation.

## 1 Results

### 1.1 SG nucleation requires mitochondrial transcripts

Stress granules (SGs) sequester diverse RNAs, including non-coding RNAs (*3*) (*4*), circRNAs, mt-tRNAs, mt-rRNAs (*4*), and a substantial set of mitochondrial protein-coding transcripts (*3, 20, 21*). Notably, mitochondrial RNAs rank among the top 25 transcripts within SGs (*3*). To validate their presence, we focused on ND5—the most abundant and smallest mitochondrial transcript reported in SGs (*3*). smFISH detection of ND5 RNA colocalized with mitochondria marked by anti-COX IV (fig. S1A). We used IMT1 to inhibit mitochondrial RNA polymerase, which reduced different mitochondrial transcripts (fig. S1F). Using smFISH, we found IMT1 specifically reduced ND5 signal in mitochondria (fig. S1, B and C), confirming probe specificity. We next examined ND5 localization during oxidative stress induced by 30-minute sodium arsenite treatment, alongside immunodetection of endogenous G3BP1-a key scaffold and marker of SG (Fig. 1A). ND5 transcripts clearly localized within SGs, as shown by line-profile analysis (Fig. 1B), supporting sequencing-based observations (*3, 21*). Interestingly, mitochondrial ND5 levels increased by 50 % within 30 minutes of oxidative stress (fig. S1, D to E), suggesting rapid mitochondrial transcriptional responses concurrent with SG formation.

**Figure 1:**
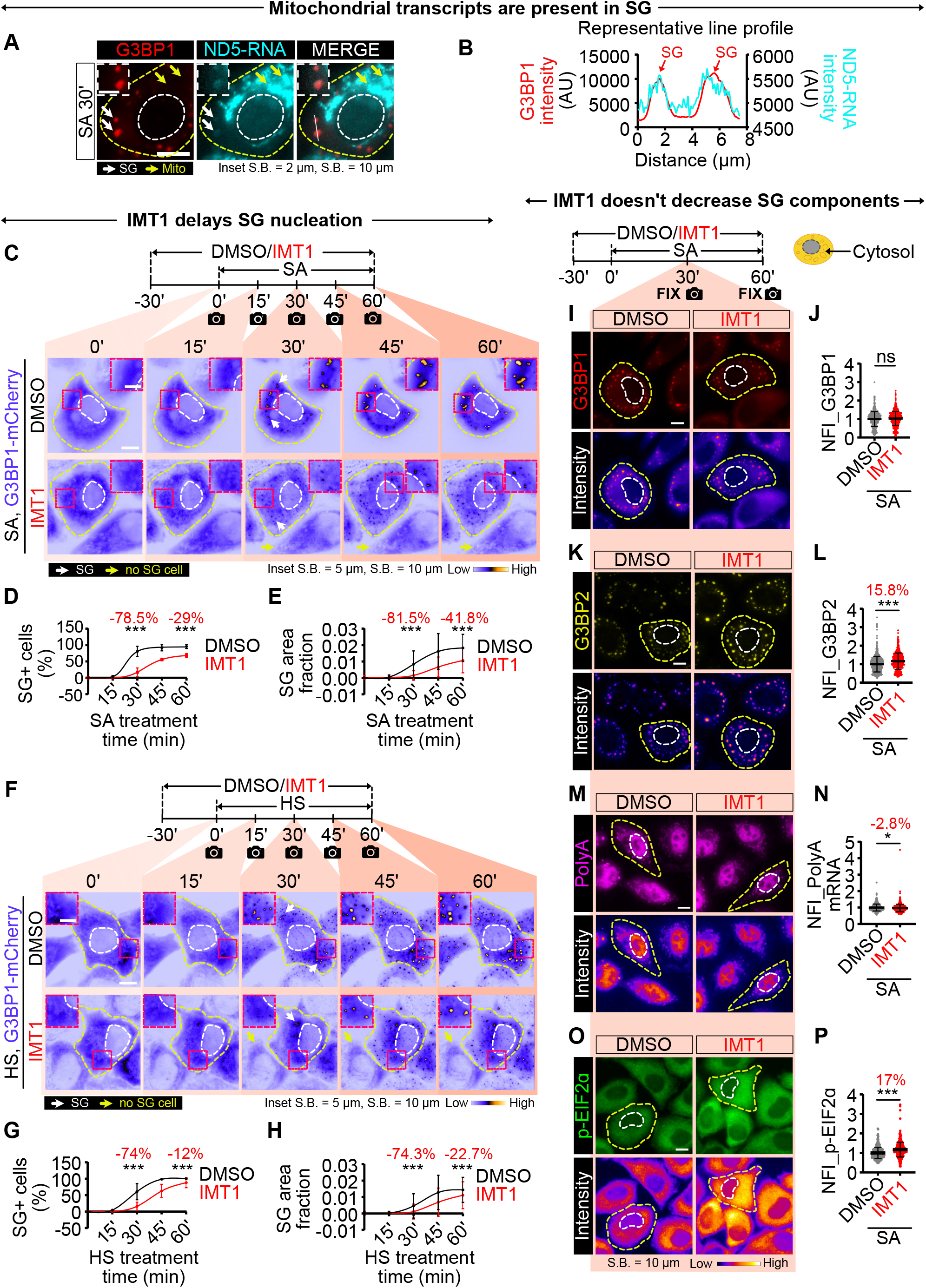
Mitochondrial transcripts are required for SG nucleation. (**A**) Immunofluorescence images of HeLa cells stained for G3BP1 (red) and mitochondrial ND5 RNA (cyan) after 30 min of oxidative stress (SA) at 37°C; yellow arrows mark mitochondrial ND5 RNA. (**B**) Fluorescence intensity profiles of ND5 and G3BP1 measured along the region indicated in (A). (**C**) Time-lapse, intensity color-coded G3BP1-mCherry images of DMSO- or mitochondrial transcription inhibitor-IMT1-treated HeLa cells during oxidative stress (t = 0 marks addition of NaAsO_2_ or SA). (**D, E**) Per cell fraction of SG-positive cells and total SG area relative to cytoplasm during oxidative stress. (**F**) Intensity color-coded G3BP1-mCherry images during 42°C-heat stress after DMSO or IMT1 treatment (t = 0 marks heat induction). (**G, H**) Per cell fraction of SG-positive cells and SG area during heat stress. (**I, K**) Immunofluorescence images of G3BP1 and G3BP2 on DMSO vs IMT1 treatment under oxidative stress; (**J. L**) Corresponding Normalised mean Fluorescence Intensity quantification for cytosol. (**M**) PolyA-tailed mRNA detected by oligo(dT) FISH in DMSO- and IMT1-treated cells after oxidative stress. (**N**) corresponding Normalised mean Fluorescence Intensity quantification for cytosol. (**O**) Immunofluorescence of p-eIF2μ after 30 min oxidative stress in DMSO vs. IMT1. (**P**) cytosolic intensity quantification. More than 360 cells were analyzed per condition across three biological replicates. Cell and nuclear outlines are shown in yellow and white, respectively. White arrows indicate SGs. Statistical significance was determined by Mann–Whitney U test (*P *<* 0.05, **P *<* 0.01, ***P *<* 0.001, ns P *>* 0.05). Percentage changes relative to the respective control are shown on the graphs.

To test whether mitochondrial transcripts contribute to SG formation, we inhibited mitochondrial transcription with IMT1 and performed live-cell imaging in HeLa cells stably expressing mCherry-G3BP1 (*22*). To avoid imaging-induced shrinkage seen at 37 °C (fig. S2A, Movie 3), SG nucleation was monitored during oxidative stress at 25 °C, with imaging every 15 minutes for 60 minutes (Fig. 1C, Movie 1). After 30 minutes of stress, only 17 % of IMT1-treated cells formed SGs compared with 80 % of controls (Fig. 1D), reflecting a 78.5 % reduction in SG+ cells. Although this difference narrowed with prolonged stress, IMT1-treated cells consistently showed fewer SG+ cells even at 60 minutes. To quantify SG formation per cell, we measured the SG area fraction (SG area normalized to cytoplasmic area), which was reduced by 81.5 % at 30 minutes and 41.8 % at 60 minutes in IMT1-treated cells (Fig. 1E). The number of SGs per cell and G3BP1 enrichment were also markedly lower with IMT1 treatment (fig. S2, F and G). Similar reductions were observed when oxidative stress was applied at 37 °C (fig. S2, A to E), indicating that the effect of mitochondrial transcription inhibition on SG nucleation is independent of imaging conditions. Together, these results show that mitochondrial transcription is critical for early SG nucleation during oxidative stress. Although SG properties partially recover by 60 minutes, mitochondrial transcripts are essential to initiate robust SG assembly.

We next asked whether mitochondrial transcripts influence SG nucleation in other stresses, such as heat shock at 42 °C((*23*), Fig. 1F, Movie 2). Similar to oxidative stress, IMT1 treatment alongwith heat stress reduced SG+ cells by 74 % at 30 minutes, and by 60 minutes IMT1-treated cells had only 12 % less SG+ cells than controls (Fig. 1G). IMT1 also decreased SG area fraction (Fig. 1H), SG number per cell (fig. S2H), and G3BP1 enrichment (fig. S2I) during heat stress. These results indicate that mitochondrial transcripts support SG nucleation across diverse stress conditions.

Because SGs form through multivalent interactions among scaffold RBPs (G3BP1, G3BP2) and free mRNAs (*22, 24, 25*), we tested whether mitochondrial transcription inhibition alters these scaffolds. IMT1 alone increased G3BP1 by 20% (fig. S3, A and B), while G3BP2 and poly(A) mRNA remained unchanged (fig. S3, C to F). Under oxidative stress, IMT1 did not reduce G3BP1 (Fig. 1I and J), increased G3BP2 by 15.8% (Fig. 1, K and L), and caused only a minor 2.8% decrease in poly(A) RNA (Fig. 1M and N). Since RNA release through translation inhibition is central to SG nucleation (*25*), we examined EIF2α phosphorylation a major indicator for cap dependent translation inhibition. IMT1 along with SA increased p-EIF2α by 17% (Fig. 1, O and P) without altering total EIF2α (fig. S3, G and H). Together, these findings show that IMT1 does not deplete SG scaffold components, suggesting that mitochondrial transcripts might play a direct role in promoting SG nucleation.

### 1.2 Mitochondrial dsRNA is required for SG nucleation

Mitochondrial DNA is transcribed bidirectionally which produces overlapping transcripts capable of forming dsRNA. Mitochondrial dsRNA increases during mitosis (*26*), neurodegeneration (*27*), ageing (*28*), and autoimmune diseases (*16, 29, 30*). Our previous in vitro and in-cellulo studies showed that ssRNA dissolves condensates, whereas dsRNA nucleates them (*18*); SGs are also enriched in endogenous dsRNA (*18*). This prompted us to ask whether mitochondrial dsRNA contributes to SG nucleation.

First, we ask if cellular dsRNA increases during stress. Using the well characterised J2 antibody which detects dsRNA (*16*), (*26*), we found 30 minutes of oxidative stress increased cytosolic dsRNA by ∼37 % (Fig. 2, A and B), while 60 minutes caused a 20 % increase (fig. S4, A and B). Heat shock similarly elevated cytosolic dsRNA by ∼ 25 % at 30 minutes and ∼13 % at 60 minutes (fig. S4, C and D). Thus, endogenous dsRNA rises early during stress.

**Figure 2:**
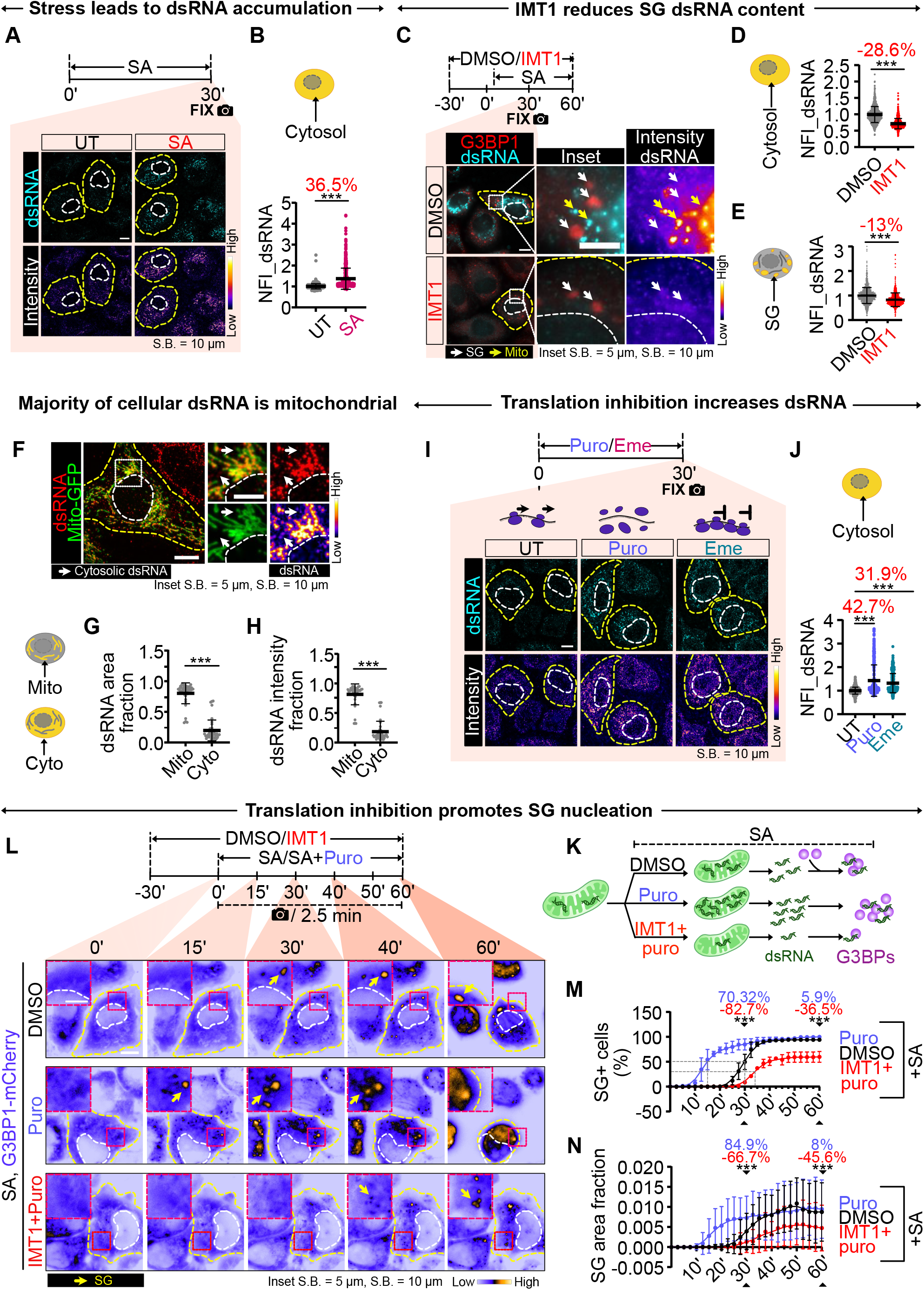
Mitochondrial dsRNA is required for SG nucleation. (**A**) Immunofluorescence of endogenous dsRNA (J2) in HeLa after 30 min oxidative stress. (**B**) Cytosolic Normalised mean Fluorescence Intensity of dsRNA across treatments (*>*600 cells/condition across three replicates). (**C**) Endogenous G3BP1 and dsRNA under DMSO- or mitochondrial transcription inhibitor-IMT1 treatment during oxidative stress. (**D, E**) Cytosolic and SG-localized dsRNA Normalised mean Fluorescence Intensity quantification. (**F**) Immunolocalization of endogenous dsRNA foci (J2) in cells expressing mito-GFP. (**G, H**) Per cell cytoplasmic dsRNA foci area fraction and fraction of cytoplasmic integrated dsRNA intensity inside mitochondria or cytosol (*>*40 mito-GFP cells). (**I**) Immunolocalisation of dsRNA in cells treated with translation inhibitors (puromycin-Puro, emetine-Eme). (**J**) Cytosolic Normalised mean Fluorescence Intensity of dsRNA across treatments. (**K**) Schematic of SG nucleation under oxidative stress in cells treated with SA alone (intermediate dsRNA), or along with Puro (highest dsRNA) and Puro+IMT1 (low dsRNA). (**L**) Live-cell imaging of G3BP1-mCherry in oxidative stress treated cells with DMSO or Puro or Puro combined with IMT1; dashed boxes denote insets; yellow arrows mark SGs. (**M**) Fraction of SG-positive cells over time calculated for at least 300 cells per condition across 3 biological replicates. Grey dotted lines shows the time at which 50 % of cells have SG for a given condition (**N**) SG area fraction over time (*>*130 cells/condition across three replicates). Data are mean ± s.d. Cell and nuclear outlines are yellow and white. Significance: Mann–Whitney U test (*P *<* 0.05, **P *<* 0.01, ***P *<* 0.001, ns P *>* 0.05). Percentage changes relative to the respective control are shown on the graphs.

To test whether reduced SG nucleation during IMT1 treatment correlates with decreased dsRNA levels, we quantified dsRNA under stress with or without IMT1 exposure (Fig. 2C). IMT1 lowered cytosolic dsRNA by ∼29 % at 30 minutes (Fig. 2D) and 47 % at 60 minutes (fig. S4, I and J). SG-localized dsRNA also fell by 13 % at 30 minutes (Fig. 2E) and 40 % at 60 minutes (fig. S4, I and K). These results indicate that SGs continuously acquire mitochondrial dsRNA during stress, which diminishes under IMT1. Importantly, IMT1 exposure at physiological conditions also reduces dsRNA, reinforcing mitochondria as the dominant dsRNA source both in stressed and unstressed conditions (fig. S4, E to K).

We next examined dsRNA localisation relative to mitochondria. Consistent with earlier studies (*16*), mito-GFP co-staining revealed that approx. 80 % of cytoplasmic dsRNA foci and total cytosolic dsRNA signal resided within mitochondria (Fig. 2, F to H) in physiological condition, confirming mitochondrial dsRNA as the major pool.

Both oxidative and heat stress—which induce SGs—also elevated mitochondrial dsRNA. Because translation inhibition occurs in these stresses and is required for SG induction (*31, 32*), we hypothesised that translation shutdown drives dsRNA accumulation. Measuring translation via puromycin incorporation indeed showed translation inhibition during oxidative stress and heat shock at both 30 minutes and 60 minutes (fig. S5, A to D). To test whether translation inhibition alone increases dsRNA, we used puromycin (polysome dissociation) and emetin (ribosome stalling). Both inhibitors inhibited translation (fig. S5, E to I) and increased cytosolic dsRNA by *>*30 % (Fig. 2, I and J), demonstrating that translation inhibition is sufficient to elevate cytosolic dsRNA.

Because translation inhibition develops progressively during stress (*33*), we reasoned that accelerating inhibition should rapidly increase dsRNA and thereby hasten SG nucleation. Indeed, combined oxidative stress (SA) and puromycin treatment during live cell imaging, triggered SG formation in 50 % of cells (shown by dotted grey lines, Fig. 2 L and M, Movie 4) within 15 minutes, compared with ∼30 minutes required for SA alone (Fig. 2L and N, Movie 4). To distinguish contributions of mitochondrial dsRNA from free mRNA released by puromycin, we added IMT1. IMT1 delayed SG nucleation relative to SA+puro and SA alone, increasing nucleation time to 34 minutes for 50 % of cells (Fig. 2M). IMT1 also decreased the maximal SG+ cells at 1 hour to ∼ 60 %, versus 100 % (SA+puro) and 94 % (SA). The SG area fraction was similarly reduced in IMT1 treatment (Fig. 2N). These results establish that stress-induced translation inhibition elevates mitochondrial-derived dsRNA, which directly accelerates SG nucleation, rather than mRNA released from ribosomes.

### 1.3 Translation stress-activated PKR remodels mitochondria and releases dsRNA

So far, we have shown that mitochondrially transcribed dsRNA is required for SG nucleation during translation inhibition. As ∼80 % of cellular dsRNA is mitochondrial (Fig. 2, F to H), but SGs form in the cytosol, we asked whether cytosolic dsRNA increases during stress. Treatment of HeLa cells with puromycin (Fig. 3, A to C) or oxidative stress (fig. S6, A to C), followed by dsRNA (J2) and TOM20 staining, revealed that cytosolic J2 intensity increased by ∼20 % (Fig. 3, A and C) and ∼12 % (fig. S6, A and C), respectively. Mitochondrial J2 foci intensity also rose by 114 % in puromycin (Fig. 3, A and B) and 33 % in SA treatment (fig. S6, A and B). These results suggest that stress triggers a mechanism releasing mitochondrial dsRNA into the cytosol.

**Figure 3:**
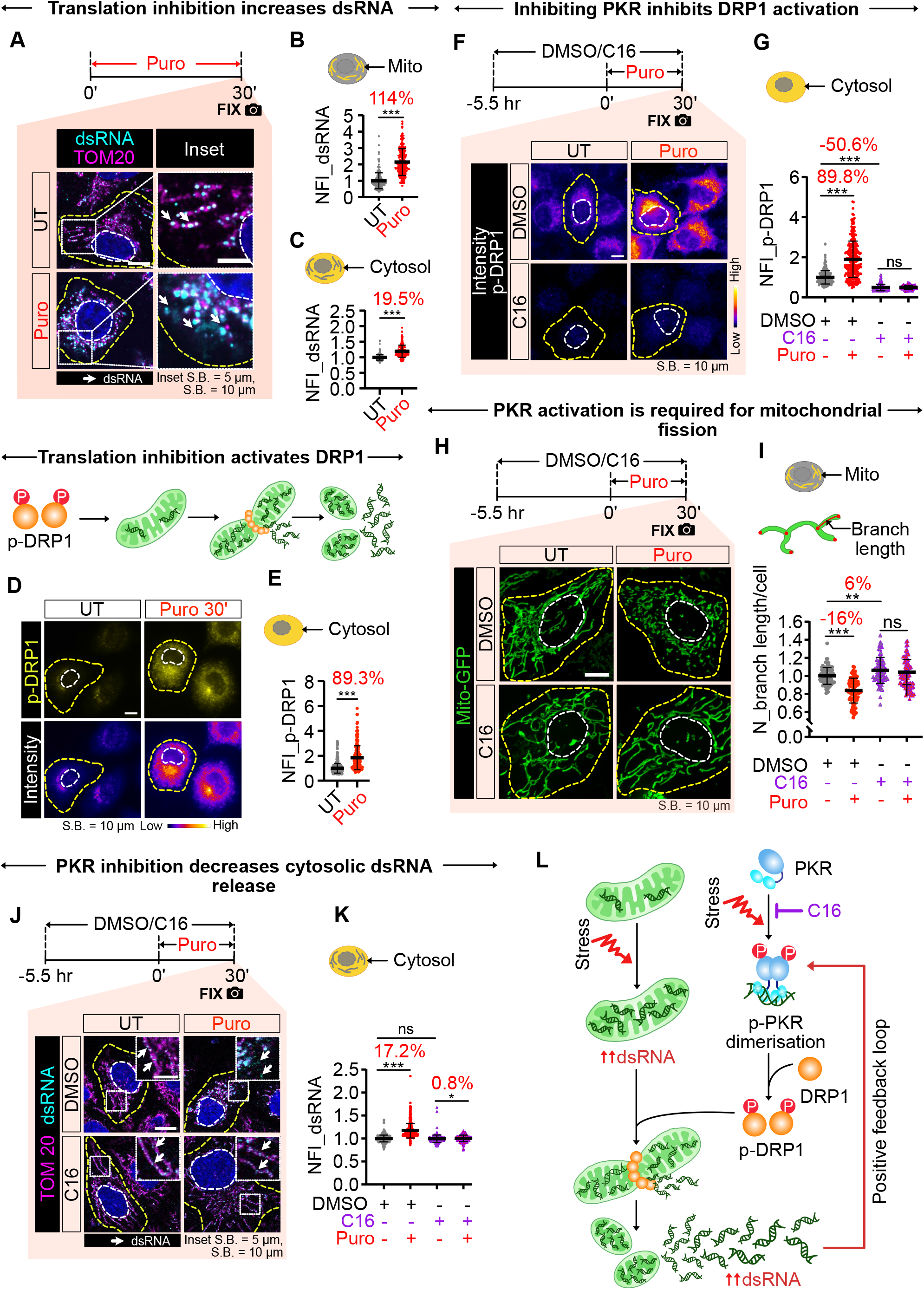
Translation inhibition triggers PKR-dependent mitochondrial fragmentation and dsRNA export. (**A**) HeLa cells stained for TOM20 (mitochondria) and dsRNA (J2); white arrows indicate dsRNA. (**B**) Mitochondrial, (**C**) Cytosolic Normalised mean Fluorescence Intensity of dsRNA. (*>*340 cells per condition across three replicates). (**D**) Immunofluorescence images showing p-DRP1 in control and Puro-treated cells. (**E**) Cytosolic Normalised p-DRP1 mean Fluorescence intensity (*>*325 cells per condition across three replicates). (**F**) p-DRP1 levels in HeLa with Puro and/or C16 (PKR inhibitor) treatment. (**G**) Cytosolic Normalised p-DRP1 mean Fluorescence Intensity across treatments (*>*160 cells per condition across three replicates). (**H**) Fluorescence images of mito-GFP expressing cells with Puro and/or C16. (**I**) Normalized mean branch length of mitochondria per cell (*>*90 cells per condition across three replicates). (**J**) Immunofluorescence representative images of TOM20 and dsRNA in cells treated with Puro and/or C16; white arrows indicate dsRNA. (**K**) Cytosolic Normalised mean Fluorescence Intensity of dsRNA. (*>*191 cells per condition across three replicates). (**L**) Model: stress elevates mitochondrial dsRNA and activates PKR, which phosphorylates DRP1 to promote mitochondrial fission; fission releases mitochondrial dsRNA to the cytosol, capable of activating PKR and forming a positive feedback loop. Data are mean ± s.d. Cell and nuclear outlines are yellow and white. Significance: Mann–Whitney U test (*P *<* 0.05, **P *<* 0.01, ***P *<* 0.001, ns P *>* 0.05). Percentage changes relative to the respective control are shown on the graphs.

A hint of how dsRNA can be released from mitochondria, came from TOM20 images which showed fragmented mitochondria during translation inhibition (Fig. 3A). To quantify this, we imaged mito-GFP–labeled mitochondria in 3D and analyzed them using a published image analysis pipeline (methods). The classifier distinguished between elongated, fragmented, and physiological states of mitochondria (fig. S7). After 30 min puromycin treatment (fig. S8A), mean branch length (fig. S8B), surface area (fig. S8D), and volume (fig. S8E) all decreased, while sphericity increased (fig. S8C), indicating a shift to smaller, more spherical mitochondria. Emetin caused similar mitochondrial fragmentation (fig. S8, F to J). Oxidative stress (fig. S9, A to J) and heat shock (fig. S9, K to T) also induced mitochondrial fragmentation at both 30 and 60 min. Thus, stress-induced translation inhibition promotes mitochondrial fragmentation.

Because fragmentation coincides with increased cytosolic dsRNA, we asked whether mitochondrial fission enables dsRNA release. Using ωDRP1 HeLa cells—lacking the core fission regulator DRP1 (fig. S10A, (*34*))—we confirmed that DRP1 KO produces elongated mitochondria (fig. S7A) that fail to remodel upon puromycin treatment (fig. S10, B to F). Consistent with our hypothesis, cytosolic dsRNA did not increase in ωDRP1 cells during puromycin treatment (fig. S10, G and H). These results indicate that mitochondrial fragmentation is required for dsRNA release during stress-induced translation inhibition.

Mitochondrial fission requires phosphorylated DRP1, which assembles into a multimeric ring around mitochondria for its constriction (*35*). We therefore tested whether DRP1 becomes phosphorylated during translation inhibition. Both puromycin (30 min; Fig. 3, D and E) and oxidative stress (fig. S11, C and D) increased p-DRP1 by ∼89 % and ∼75 %, respectively. This was not due to elevated DRP1 abundance as oxidative stress reduced total DRP1 (fig. S11, A and B). Stress-associated kinases are known to phosphorylate DRP1 (*36, 37*). Because oxidative stress also activates the dsRNA-sensing kinase PKR (*38*), we hypothesized that PKR activates DRP1. Indeed, PKR became phosphorylated during oxidative stress and puromycin treatment (fig. S11, E to G). PKR activation requires phosphorylation, often utilising the inorganic phosphate released by ATP bound to the PKR ATP-binding pocket (*39*). We used C16, an inhibitor of the PKR ATP-binding pocket (*40*), effectively blocked p-PKR induction on poly (I:C) treatment, a known PKR activator (fig. S12, A to C) (*41*). C16 co-treatment with puromycin reduced p-DRP1 levels (Fig. 3, F and G) and rescued mitochondrial fission, as branch length (Fig. 3, H and I), surface area (fig. S12F), volume (fig. S12G), and sphericity (fig. S12E) remained unchanged relative to controls. C16 did not alter total DRP1 abundance under any condition (fig. S13, B and C). Interestingly, C16 also decreased the physiological p-DRP1 levels (Fig. 3, F and G), resulting in elongated mitochondria in physiological conditions as well (Fig. 3 H and I). Inhibiting mitochondrial fragmentation via C16 treatment also abolished the increase in cytosolic dsRNA (Fig. 3, J and K).

Collectively, these results show that stress or translation inhibition elevates cytosolic dsRNA, activates the dsRNA-sensing kinase PKR, and drives PKR-dependent DRP1 phosphorylation to remodel mitochondria (Fig. 3 L). Thus, PKR-mediated control of DRP1 not only enables stress-induced mitochondrial fragmentation but is also essential for maintaining mitochondrial morphology during physiological homeostasis.

### 1.4 SGs nucleate at the mitochondria-ER contact sites and buffer inflammation

In previous sections we showed that stress and translation inhibition increase mitochondrial dsRNA and promote SG nucleation. Mitochondrial dsRNA is a strong inflammatory trigger (*16, 27*), consistent with the rapid induction of inflammatory cytokine transcripts (fig. S14A) and p-IRF3, an inflammatory transcription factor (fig. S14, B and C), upon translation inhibition. As SGs buffer dsRNA-induced inflammation (*18*), we asked whether SGs and SG-scaffold RBPs mitigate inflammation under translation inhibition. WT U2OS cells, and complete G3BP1/2 KO (ΔG3BPs) cells (*42*) were treated with puromycin, and inflammatory cytokines were quantified (fig. S14D). Loss of SG scaffolds led to markedly higher inflammation than WT, indicating that SG-RBPs help buffer dsRNA-driven inflammation, although this does not exclude direct roles of G3BPs independent of SGs.

To test the contribution of SG condensates specifically, we used optogenetically induced SGs formed by mCherry-G3BP1 with Cry2 (opto-G3BP1) (*43*) and quantified p-IRF3 as a readout of inflammation (Fig. 4A). mCherry-G3BP1 lacking Cry2 served as a control and did not form optoSGs on photoactivation (Fig. 4B; fig. S14, E and F). G3BP1 overexpression alone reduced p-IRF3 levels, as seen from intensity-coded images and scatter plots (Fig. 4, B and C-left), where p-IRF3 and G3BP1 levels were inversely correlated. Formation of optoSGs further decreased p-IRF3 compared to similar G3BP1 levels (Fig. 4C-right), evident from a approximately two-fold steeper slope for opto-G3BP1 compared with G3BP1 (Fig. 4C-left). Thus, condensates buffer inflammation more effectively than scaffold proteins alone.

**Figure 4:**
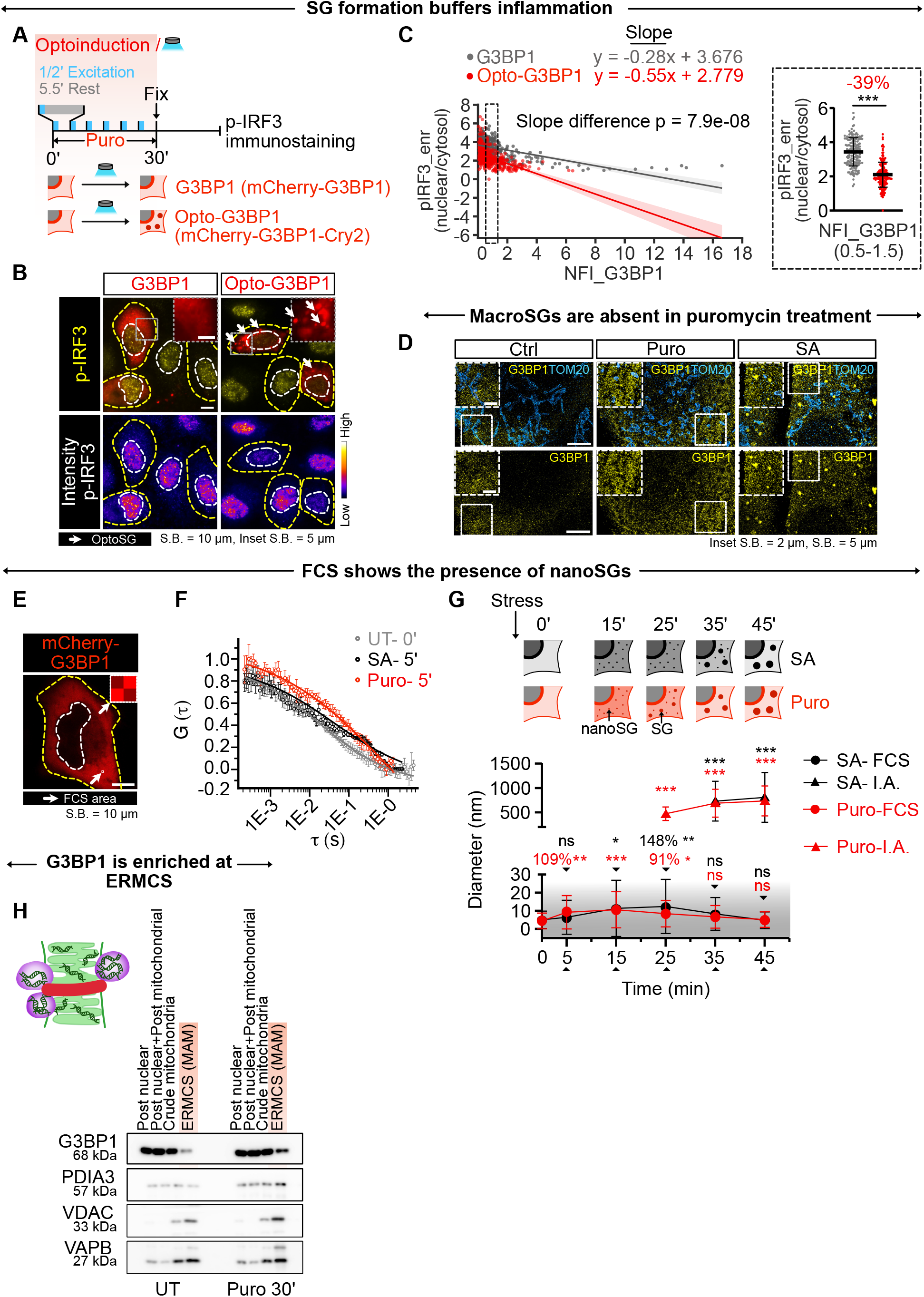
SG nucleate at mitochondria-ER contact sites and buffer inflammation. (**A**) Schematic of optogenetic activation of G3BP1 or opto-G3BP1 and downstream inflammation monitoring by p-IRF3 quantification. (**B**) Representative immunostaining of p-IRF3 in cells expressing G3BP1 variants after photoactivation. (**C**) Left-scatter plot of Normalised mean Fluorescence intensity of mCherry–G3BP1 levels versus nuclear p-IRF3 enrichment in optogenetically activated G3BP1 variants. The solid lines represent linear regression with 95 % confidence represented by the shaded region. The P value of the slope difference between linear regressions of G3BP1 variants was calculated using the ANCOVA test. Right graph showing the p-IRF3 enrichment values selected for Normalized mean Fluorescent Intensities of G3BP1 between 0.5-1.5. (*>*X cells across three replicates). (**D**) DNA-PAINT super-resolution images of HeLa cells treated for 30 min with puromycin or oxidative stress, stained for mitochondria (TOM20) and SGs (G3BP1). (**E**) HeLa cell expressing mCherry-G3BP1 image with region of interest (white box) for FCS measurement. (**F**) Normalized autocorrelation function G (τ) from point FCS measured in cytoplasm without stress (UT–0’) and 5 min after oxidative stress (SA-5’) or translation inhibition treatment (Puro–5’). (**G**) The grey region of the graph shows the diameter of nanoSG as a function of treatment time, derived from FCS diffusion coeffcients and microviscosity measurements. Quantification was done for at least 24 FCS measurements per condition over three biological replicates. The % change and p values for the difference are shown with respect to the 0 time point of the corresponding treatment on the graph. Refer to methods for p value calculations. The white region of the graph shows the diameter of macro SGs as computed from confocal images using Image Analysis Fiji ImageJ (I.A.). (**H**) Immunoblots of subcellular fractions showing stress granule marker-G3BP1, ER lumen protein-PDIA3, mitochondrial outer membrane protein -VDAC, and ER-MC contact protein VAPB, untreated versus puromycin-treated cells. Data are mean ± s.d. Cell and nuclear outlines are yellow and white. Significance: Mann–Whitney U test (*P *<* 0.05, **P *<* 0.01, ***P *<* 0.001, ns P *>* 0.05), unless specified otherwise. Percentage changes relative to the respective control are shown on the graphs.

Despite this buffering, endogenous G3BP1 formed no large SGs after puromycin treatment (Fig. 4D), consistent with earlier reports (*25,44*). DNA-PAINT super-resolution imaging again showed smaller mitochondria in puromycin-treated cells (Fig. 4 D, fig. S17C). Under oxidative stress, SGs were visible, but under puromycin we observed only clustered G3BP1 signals (Fig. 4D). These observations suggested that during translation inhibition, G3BP1 forms sub-resolution nanocondensates or nanoSGs that buffer inflammation.

To detect nanoSGs, we used Fluorescence Correlation Spectroscopy (FCS) to measure mCherry-G3BP1 diffusion through confocal volumes (Fig. 4E). Slow-moving G3BP1 species shifted the autocorrelation function G(τ) to the right (Fig. 4F) which was fitted to extract the diffusion coefficient ((D), methods). By applying the Stokes-Einstein relation, we inferred condensate diameter from the estimated cytoplasmic viscosity (η) (fig. S15, A–J) and D (fig. S16B).

NanoSGs formed within 5 minutes of induction specifically: puromycin generated 9 ± 9 nm nanoSGs, whereas SA produced nanoSGs which were very similar to the untreated samples (6.4 ± 9.4 nm diameter) (Fig. 4G), indicating faster nucleation upon translation inhibition. NanoSGs grew more rapidly in puromycin treatment, reaching 10.5 ± 10 nm by 15 min, whereas SA reached 12.4 ± 14.9 nm at 25 min, consistent with live-cell SG nucleation assays. At 25 min, puromycin—but not SA—showed both nanoSGs and emerging ∼500 nm macro-condensates (Fig. 4G-top graph, fig. S16A). By 35 min, both stresses formed ∼700 nm macroSGs, accompanied by a decline in nanoSG diameter (Fig. 4G-top graph, fig. S16A). NanoSGs never exceeded 30 nm in any of our measurements, suggesting that sub-30 nm condensates grow into macroSGs, after which the cytoplasm becomes depleted of nanoSGs, resembling unstressed cells by 45 min (Fig. 4G). Overall, puromycin is a stronger inducer of both nano- and macroSGs, and macroSG formation restores cytoplasmic homeostasis by removing immunogenic dsRNA.

To test whether dsRNA released during mitochondrial fragmentation drives nanoSG formation, we performed FCS in HeLa ωDRP1 cells expressing mCherry-G3BP1 (fig. S16C). Loss of DRP1 prevents puromycin-induced mitochondrial fragmentation (fig. S10, B to F), cytosolic dsRNA release (fig. S10, G and H), and delays SG nucleation (fig. S16E). NanoSGs were consistently smaller in ΔDRP1 cells across all time points (fig. S16E) and reached diameter of only 8 ± 11.9 nm at 15 min. No large (*>*200 nm) SGs formed even after 1 h (data not shown). Thus, in ΔDRP1 cells nanoSGs fail to grow due to the absence of cytosolic dsRNA released by mitochondrial fragmentation. We conclude that nanoSGs form immediately upon dsRNA release from fragmented mitochondria, grow over time, and are required for macroSG formation.

p-DRP1 is required for ER–mitochondria contact site (ERMCS) formation and mitochondrial constriction (*45, 46*). As nanoSG formation depends on dsRNA generated by DRP1-mediated mitochondrial fission, we hypothesized that ERMCSs may serve as the sites of dsRNA release for nanoSG nucleation. Notably, large fraction of oxidative-stress-induced SGs localize near ERMCSs, and G3BP1 is enriched at these sites (*47*). ERMCSs were isolated after 30 min puromycin (Fig. 4H) and 30 and 60 min SA (fig. S17, A and B) using established protocols (*47*). Enrichment of VAPB (ER) and VDAC (mitochondrial) confirmed successful isolation of ERMCSs (Fig. 4H). G3BP1 was more enriched at ERMCSs after 30 min puromycin (Fig. 4H) than after SA (fig. S17, A and B), supporting the idea that translation inhibition triggers stronger dsRNA release and SG nucleation. Longer stress reduced G3BP1 enrichment at ERMCSs (fig. S17, A and B), suggesting that SGs nucleate there but subsequently move away. The absence of macroSGs coincided with pronounced mitochondrial fragmentation under puromycin treatment. In contrast, under oxidative stress, which induces macroSG formation, mitochondria were less fragmented than under puromycin treatment but more fragmented than under control conditions. These observations suggest that macroSGs might play an important role in maintaining mitochondrial morphology and integrity (fig. S17C).

Together, these results show that nanoSGs nucleate at ERMCSs, the likely sites of dsRNA release during stress-induced mitochondrial fragmentation. These nanoSGs buffer inflammation and mature into macroSGs, restoring cytoplasmic homeostasis by clearing immunogenic dsRNA from the cytosol.

## 2 Discussion

Cells organize the cytosolic milieu through membrane-bound organelles and membrane-less condensates. Cytosolic RNA–RBP condensates—stress granules (SGs)—interact with organelles to maintain organellar integrity (*5, 7*). Mitochondria, which are structurally and metabolically dynamic, physically associate with SGs (*8*), and SGs contain mitochondrial transcripts (*3, 4, 20, 21*). However, the mechanistic relationship between mitochondria and SGs has remained unclear. Our results show that stress-induced mitochondrial fragmentation releases mitochondrial dsRNA at ERMCSs, which rapidly nucleates SGs and may prevent further mitochondrial fragmentation and inflammation (Fig. 5).

**Figure 5:**
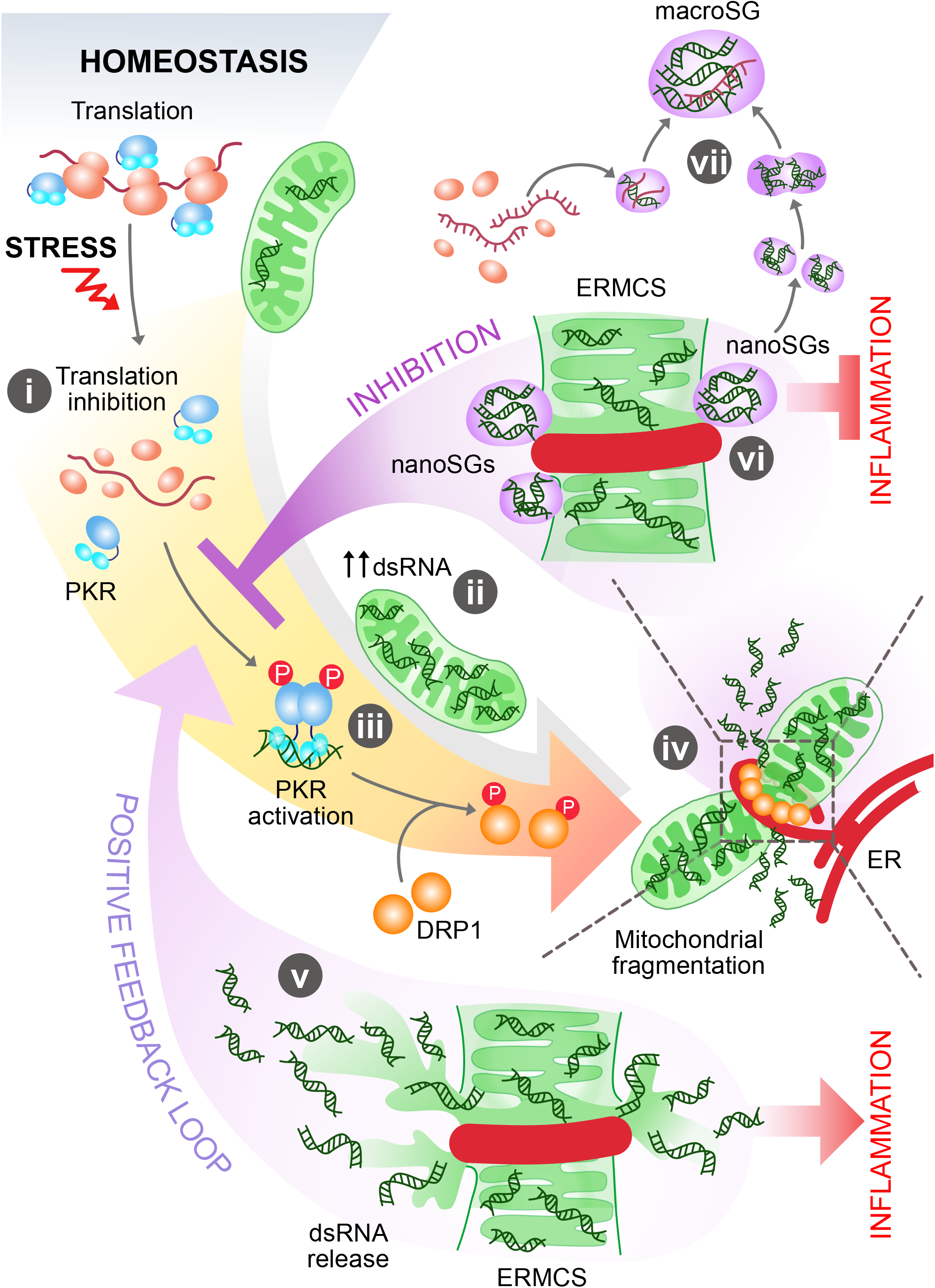
Schematic illustrating how stress granules (SGs) buffer inflammation by limiting dsRNA-driven mitochondrial remodeling. (**1**) Diverse cellular stressors inhibit translation, directly activating PKR. (**2**) In parallel, translation inhibition elevates mitochondrial double-stranded RNA (dsRNA) levels. (**3**) Activated PKR phosphorylates the mitochondrial fission factor DRP1. (**4**) Phosphorylated DRP1 assembles into constriction rings around mitochondria, promoting ER–mitochondria contact site (ERMCS) formation and driving mitochondrial fission. (**5**) Fission at ERMCSs releases mitochondrial dsRNA into the cytosol, amplifying PKR activation and triggering extensive mitochondrial fragmentation, while also inducing innate inflammatory signaling. (**6**) The dsRNA released at ERMCSs nucleates G3BP1 to form nano–stress granules (nanoSGs), which sequester dsRNA and occlude ERMCSs to restrict further mitochondrial dsRNA release. This buffering action suppresses inflammation and interrupts the positive feedback loop that would otherwise intensify mitochondrial fragmentation. (**7**) NanoSGs subsequently expand to form macroSGs either by fusing with one another or by accumulating additional mRNAs and RNA-binding proteins.

Translation inhibition—a common downstream consequence of many stresses (*33*) (Fig. 5)—is sufficient to induce dsRNA-mediated inflammation by Fig. 5(i) increasing mitochondrial dsRNA and Fig. 5(ii) activating PKR via phosphorylation. Activated PKR drives DRP1-dependent Fig. 5(iii) mitochondrial fission Fig. 5(iv), a leaky process that releases additional dsRNA, establishing a positive feedback loop Fig. 5(v) that exacerbates PKR activation and mitochondrial damage. This loop is curtailed by rapid dsRNA-driven nucleation of nanoSGs at ERMCSs Fig. 5(vi). Further the nanoSGs can grow to form macroSGs Fig. 5(vii).

Mitochondrial transcription is essential for SG nucleation across stresses, consistent with the detection of mitochondrial transcripts in SGs by sequencing (*3, 4, 20, 21*) and smFISH (Fig. 1, A and B). Short IMT1 treatment (30 minutes) did not alter SG scaffold protein or mRNA levels (Fig. 1, I to N) but reduced both cytosolic and SG dsRNA, correlating with decreased SG formation in live cell imaging (Fig. 2). While prior studies showed that dsRNA promotes condensate nucleation (*18*), our data specifically demonstrate that mitochondrial dsRNA nucleates SGs (Fig. 1, C to H) which explains the prescence of mitochondrial transcripts in SGs. Live cell imaging visualizes macroSGs (*>*200–300 nm, appearing after 25 min), but visualization of nucleation kinetics using FCS, detected 5–10 nm nanoSGs forming as early as 2–5 min after translation inhibition. NanoSGs grow to 30 nm before forming macroSGs (Fig. 4G) either by fusion or by recruiting additional mRNAs and RBPs (Fig. 5 (v)), explaining why nanoSGs larger than 30 nm are absent once macroSGs form. As macroSGs mature (25–45 min), cytosolic nanoSG levels progressively decline, and by 45 min the cytosol resembles the unstressed state (Fig. 4G). High-resolution FCS showed that direct translation inhibition nucleates nanoSGs faster than oxidative stress, reinforcing that translation inhibition alone is sufficient for SG nucleation. Notably, DRP1-deficient cells—unable to release mitochondrial dsRNA—still form nanoSGs, likely from preexisting cytosolic dsRNA, but these do not grow beyond 8 nm, indicating that SG maturation requires ongoing mitochondrial dsRNA release. Thus, SG nucleation depends on mitochondrial dsRNA, while mRNA released from stalled ribosomes likely supports macroSG assembly.

Our findings suggest that SGs buffer inflammation primarily by regulating mitochondrial fragmentation. Mitochondria are major sources of inflammatory signals, including ROS (*48*), cyt C (*17*), dsDNA (*14*), and dsRNA (*16*). We show that mitochondrial morphology is controlled by PKR in both physiological and stress conditions (Fig. 3, H and I). Activated PKR phosphorylates DRP1 (Fig. 3, D and E), enabling DRP1 to assemble into constriction rings and remodel ER to facilitate fission. Cytosolic PKR is normally inhibited by ribosome binding, and translation inhibition activates PKR (*49*). Thus, PKR activation by stress or translation inhibition initiates mitochondrial fission and dsRNA release (Fig. 3A,B; fig. S6, A and B). Increasing cytosolic dsRNA amplifies PKR activation, reinforcing the fission loop and damaging mitochondrial structure (Fig. 5 (v)). p-DRP1 promotes ERMCSs formation (*45,46*), where VDAC may facilitate dsRNA leakage during fission (*50*). Released dsRNA rapidly nucleates nanoSGs that buffer inflammation (Fig. 4, B abd C,fig. S14D). ERMCSs serve as initial SG nucleation sites (Fig. 4H): at 30 min of oxidative stress, they are enriched for G3BP1, but by 60 min this enrichment diminishes (fig. S17A). Overexpressing G3BP1 reduces inflammation, and enhancing condensate formation via optoinduction suppresses inflammation even more effectively, likely by improving dsRNA sequestration (*18*) and/or plugging ERMCSs pores to prevent further dsRNA leakage. Both mechanisms break the PKR–fission positive feedback loop, stabilizing mitochondrial morphology. Hence, SGs buffer inflammation by controlling mitochondrial morphology.

Chronic conditions such as Huntington’s disease (*30*), osteoarthritis (*29*), diabetes (*51, 52*), neurodegeneration (*13,27,53*), and aging (*28,53*) exhibit increased dsRNA and mitochondrial fragmentation. We propose that targeting SG pathways in these diseases may offer therapeutic opportunities by restoring SG-led mitochondrial morphology, integrity, and inflammation homeostasis.

## Acknowledgments

We thank members of Dr. Shovamayee Maharana’s laboratory for their insightful discussions. We are grateful to the IISc central imaging facility, MCB departmental facility for technical support and DST-Fund for Improvement of S and T Infrastructure (FIST), University Grands Commision (UGC) Centre for Advanced Study, and DBT-IISc Partnership Program (Phase II at IISc, BT/PR27952/INF/22/212/2018) for infrastructural support. We thank Prof. Ravi Manjithaya (JNCASR, Bengaluru) and Ritopruva Sen for guidance with the mitochondrial analysis pipeline. We also thank Dr. Mainak Bose and Jeet Mukhopadhyay for assistance with single-molecule RNA-FISH probe design and labeling. We acknowledge Prof. Sandhya S. Visweswariah (IISc, Bengaluru) and Prof. Ravi Muddashetty (CBR, Bengaluru) for valuable discussions.

## Author contributions

S.M. conceived the project, designed the experimental strategy, and supervised the study. P.N. co-conceived the project, designed and carried out experiments, and performed data analysis. S.S. and J.B. conceptualized, performed and analyzed FCS experiments and helped in writing relevant parts of the manuscript. A.L. and J.J. conducted subcellular fractionation experiments for ERMCS. S.K. and N.F. carried out mitochondrial morphology analysis. P.G.D. performed dsRNA quantification under stress conditions. J.S. performed polysome profiling. A.B. and M.G. carried out super-resolution DNA-PAINT super-resolution imaging. All authors contributed to data interpretation. P.N. and S.M. wrote the manuscript.

## Data and materials availability

Further information and requests for reagents and resources should be directed to and will be fulfilled by the lead contact, Shovamayee Maharana. This study did not generate new, unique reagents.

## Materials and Methods

### Cell culture and drug treatments

HeLa cells (a gift from Dr. Sachin Kotak, Indian Institute of Science, Bengaluru, India), HeLa mCherry–G3BP1 BAC–stable cells (a gift from Prof. A. Hyman, Max Planck Institute of Molecular and Cell Biology, Dresden, Germany) (*1*), HeLa ωDRP1 cells (a gift from Prof. Mike Ryan, Monash University, Docklands, Australia), and human osteosarcoma U2OS WT and U2OS ωG3BP1/2 cells (were provided by Dr. Paul J. Anderson, Harvard Medical School and Brigham and Women’s Hospital, Boston, MA 02115) (*2*) were maintained in high–glucose DMEM supplemented with 5 % FBS and 1 % penicillin–streptomycin at 37 °C and 5 % CO_2_. Media for HeLa mCherry–G3BP1 BAC–stable cells were additionally supplemented with 5 μM blasticidin.

#### Translation inhibition

Cells were treated with 7 μg/mL puromycin ((*3*); stock in Milli-Q water) or 40 μM emetine ((*4*); stock in Milli–Q water) for the indicated times. Translation inhibitor effcacy was assessed by polysome profiling. Puromycin treatment caused a ∼3–fold increase in the monosome–to–polysome ratio (Fig. S5, E and G), whereas emetine produced no detectable shift, consistent with its ability to stall elongating ribosomes (Fig. S5, F and G). To quantify its effiect on translation rates, we performed puromycin–assisted nascent peptide labeling, which revealed an ∼81% reduction in translation following emetine treatment (Fig. S5, H and I).

#### Mitochondrial transcription inhibition

Cells were treated with 5 μM IMT1 ((*5, 6*); stock in DMSO) for the indicated durations. Inhibition effciency was confirmed by ND5–RNA FISH, which showed reduced mitochondrial transcript abundance (Fig. S1, B and C).

#### PKR inhibition

To inhibit PKR, cells were treated with 1 μM of the imidazolo–oxindole inhibitor C16 ((*7*); stock in DMSO) for 6 h. Effcacy was evaluated in poly(I:C)–treated cells, which robustly induce PKR phosphorylation (*8*). Poly(I:C) increased PKR phosphorylation by ∼90%, whereas C16 co–treatment reduced phosphorylation by ∼40%, indicating effective PKR inhibition (Fig. S12, A and B).

#### Mitochondrial fragmentation

Mitochondrial fragmentation was induced by treating cells with 10 μM CCCP for 1 h (*9*).

### Plasmids and transient transfections

The plasmids used in this study included pAcGFP1–Mito—referred to as mitoGFP throughout the text (a gift from Prof. Oishee Chakrabarti, Saha Institute of Nuclear Physics, HBNI, West Bengal, India)—for visualizing mitochondrial morphology; TH1343–pTT6–G3BP1–mCherry–Cry2– E490R for optogenetic induction of stress granules (a gift from Prof. A. Hyman, Max Planck Institute of Molecular and Cell Biology, Dresden, Germany); and pcDNA3.1–mCherry–G3BP1 (a gift from Prof. A. Hyman, Max Planck Institute of Molecular and Cell Biology, Dresden, Germany) for mCherry–G3BP1 expression in mammalian cells.

For live–cell imaging experiments requiring plasmid transfection, approximately 0.3 × 10^6^ HeLa cells were seeded in coverslip bottom live cell dishes (GreenFocus Research Technologies, Bengaluru, India) and cultured overnight. The following day, cells were transfected with 1 μg plasmid DNA using polyethyleneimine (PEI) mixed in serum–free DMEM. After 6 h, the transfection mix was replaced with complete media containing 5 % FBS, and cells were allowed to express the protein for an additional 12–16 h under standard culture conditions. Transfection of mitoGFP was performed using the Xfect Transfection Reagent according to the manufacturer’s instructions.

### Stress granule induction

Stress granules (SGs) were induced using three approaches: oxidative stress, heat shock, and optogenetic activation.

#### a) Oxidative stress–induced SG formation

SGs were induced by treating cells with culture medium supplemented with 0.25 mM sodium arsenite ((*10*) stock prepared in Milli–Q water) for 30–60 min at 37 ^∘^C and 5 % CO_2_ in a humidified incubator. For temperature–controlled imaging experiments, SGs were induced by applying arsenite–containing medium prepared in 10 mM HEPES buffer on a temperature–regulated microscope stage set to the desired temperature.

#### b) Heat shock–induced SG formation

For heat–shock–mediated SG induction, cells were incubated in culture medium containing 10 mM HEPES at 42^∘^C for 30–60 min as described previously (*11*) in a humidified incubator lacking CO_2_.

#### c) Optogenetic SG induction

For optogenetic induction of SGs, cells were transfected with TH1343–pTT6–G3BP1–mCherry– Cry2 E490R to express G3BP1–mCherry–Cry2. Transfection was performed using PEI in serum– free medium for 6 h, followed by recovery in complete medium containing 5 % FBS for 12– 16 h. Cry2 oligomerization was triggered using brief blue–light pulses (488 nm excitation; dwell time 5 μs; 200 mW power at source) delivered through a 40x/0.60 NA air objective on a Nikon ECLIPSE Ti2 microscope. Activation was performed using 25 % laser power applied in 30 s pulses every 5 min. These illumination parameters were optimized to robustly induce Cry2–dependent assemblies while minimizing phototoxicity and irreversible aggregation (Fig. S14, A and B). mCherry–G3BP1 oligomers were imaged immediately after activation and again at 5 min and 10 min using 561 nm excitation on a CoolLED PE–800 light source. Blue–light exposure was limited strictly to the activation windows (*12*).

### Live cell imaging of SG nucleation

SG nucleation was monitored in live HeLa mCherry–G3BP1 BAC–stable cells (*1*). Approximately 0.3 × 10^6^ cells were seeded onto No. 1, 22 mm–diameter round coverslips placed in 6–well plates and cultured overnight at 37 °C and 5 % CO_2_. The following day, after completing the required drug treatments, coverslips were transferred to a stainless–steel live–cell imaging chamber containing culture medium supplemented with 10 mM HEPES. Chambers were mounted onto a microscope equipped with an on–stage temperature–controlled incubator. SGs were induced directly on the microscope by treating cells with 0.25 mM sodium arsenite for 1 h at the indicated temperatures. No additional CO_2_ was supplied during imaging. Cells were imaged using a 60x/1.518 NA oil– immersion objective (Nikon ECLIPSE Ti2). During oxidative–stress induction at 37 °C, images were acquired every minute, but this resulted in noticeable cell shrinkage. To prevent phototoxicity and maintain cell integrity, imaging frequency was reduced to every 15 min for oxidative–stress and heat–shock experiments at 25°C (Fig. 1, C to H), and to every 2.5 min for SG nucleation dynamics on increasing dsRNA (Fig. 2, J–M). Images were acquired using the 561 nm laser line of the CoolLED PE–800 at 10 % excitation intensity with a 140 ms exposure time at the best focal plane, providing optimal signal.// Quantification of SG–positive cells was performed in Fiji using the Cell Counter plugin (*13*). SG area fraction, number of SGs per cell, and G3BP1 enrichment in SGs were quantified using a custom MATLAB script (R2024b, 24.2.0).

### Immunofluorescence

For immunofluorescence experiments, approximately 0.04x 10^6^ cells were seeded onto 12 mm– diameter No. 1 round coverslips in 24–well plates and grown overnight. The following day, cells were fixed with 4 % formaldehyde for 10 min, permeabilized with 0.25 % Triton X–100 for 10 min, and blocked for 2 h at room temperature in 2 % BSA prepared in 1x PBS. Coverslips were then incubated overnight at 4 ^∘^C with primary antibodies diluted in 2 % BSA/PBS in a humidification chamber.

After washing with 1x PBS thrice, cells were incubated for 2 h at room temperature in the dark with Alexa Fluor–conjugated secondary antibodies (1:1000 dilution in 2 % BSA/PBS). Following three additional 1x PBS washes, nuclei were stained with DAPI for 10 min. Coverslips were mounted onto glass slides using Mowiol–based mounting medium (10 % w/v Mowiol^®^ 4–88, 2.5 % DABCO, 25 % glycerol, 0.1 M Tris–Cl, pH 8.5) and sealed with transparent nail polish. Details of all primary and secondary antibodies used in this study are provided in the Supplementary Table.

### In situ RNA FISH hybridisation to detect ND5 and poly(A)–tailed mRNA

#### RNA probe design and preparation

##### a) Design of ND5 smFISH Probes

A set of ∼30 non–overlapping DNA oligonucleotides (18–22 nt each, Table 1) targeting the coding region of ND5 RNA was generated using a custom R–based design tool (written by Imre Gaspar, Max Planck Institute of Biochemistry, Munich, Germany). Probe design parameters included a GC content between 45–60 % and a minimum spacing of two nucleotides between adjacent probes. Sequences for all probes are provided in the Supplementary Table. Oligos, supplied either desalted or purified by reverse–phase cartridge, were obtained from Barcode Biosciences Pvt. Ltd., Bengaluru, India and reconstituted to 100 pmol/μL in nuclease–free water.

**Table 1:**
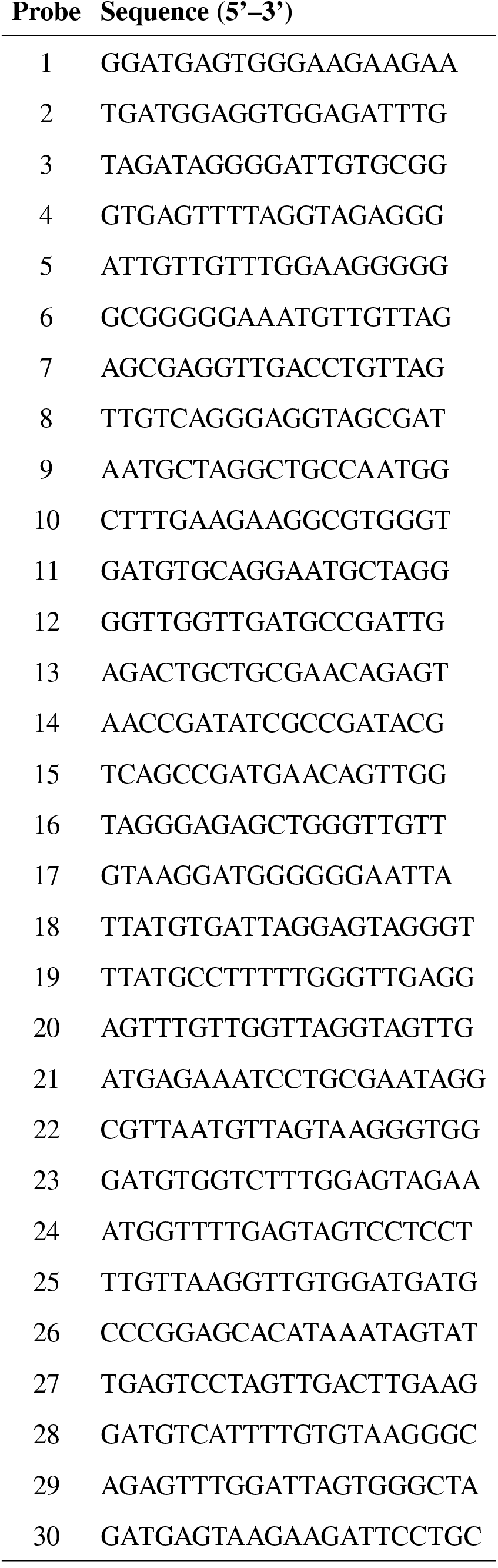
Single molecule RNA FISH probes targeting ND5.

##### b) Design of Poly(A) mRNA Probes

For detecting polyA–tailed mRNA, 3^’^–biotinylated 25–mer oligo–dT was used. Desalted or reverse– phase cartridge–purified PCR oligos were obtained from Barcode Biosciences Pvt. Ltd. Bengaluru, India.

#### 3’ end labeling of RNA probes

Two different labeling strategies were used: (i) fluorescent labeling with Atto633–ddUTP, and (ii) biotin incorporation using biotin–14–dATP. The biotin–14–dATP was detected using Streptavidin– Alexa Fluor 488.

##### a) Fluorescent 3’ end labeling using Atto633–ddUTP

i. **Synthesis of Atto633–conjugated ddUTP**: Amino–11–ddUTP (Lumiprobe #A5040) was coupled to Atto633 NHS–ester (Leica Microsystems #AD–633–31) by reacting a 1:2 molar ratio of nucleotide to dye in 0.1 M NaHCO_3_ (pH 8.3). The reaction was allowed to proceed for 2–3 h at room temperature in the dark. The residual unreacted NHS–esters were quenched using 1 M Tris–HCl (pH 7.4).
ii. **Fluorescent labeling of probe mix**: All 30 oligos were combined into a single probe pool at ∼250 μ M. Approximately 300 pmol of this mixture was labeled with 5 mM Atto633–ddUTP using the Terminal Deoxynucleotidyl Transferase (TdT) kit according to the manufacturer’s instructions.
iii. **Purification of labeled probes**: Following the TdT reaction, labeled probes were adjusted to 200 μ L with nuclease–free water containing 300 mM sodium acetate (pH 5.5) and 5 μ g linear acrylamide. Oligonucleotides were precipitated by adding 800 μ L ice–cold (–20 ^∘^C) ethanol and incubating at –80 ^∘^C for 15–20 min. Probes were pelleted by centrifugation at 16,000 g for 20 min at 4 ^∘^C, washed three times with chilled 80 % ethanol, air–dried, and finally resuspended in 25 μ L nuclease–free water.
iv. **Determination of labeling effciency**: Concentration and dye–to–probe ratios were calculated as described in Gaspar (*RNA*, 2017).

##### b) Biotin–based 3^’^ end labeling

Approximately 5 pmol of ND5 probes were labeled with 8–10 μ M biotin–14–dATP using TdT following the supplier’s instructions. Ten microliters of the completed reaction was used directly in 100 μ L of hybridization buffer.

#### Sample preparation and hybridization

For smFISH, ∼0.04 × 10^6^ cells were plated onto 12 mm No. 1 round coverslips in 24–well plates and cultured overnight. The next day, after the required treatments, cells were fixed at 37 ^∘^C using 4 % formaldehyde for 10 min. Cells were then permeabilized with 0.25 % Triton X–100 in PBS for 10 min at room temperature.

##### a) Hybridization

Coverslips were transferred to a humidified chamber and incubated in hybridization solution (125 nM labeled DNA probes, 4x SSC buffer, 0.1 M sodium citrate, 0.6 M NaCl, 5 % dextran sulfate, 10 % formamide, 0.5 mM EDTA, and 1% BSA). Samples were incubated at 37 °C for 3 h. For detection of biotin–tagged probes, coverslips were subsequently incubated with Streptavidin– Alexa Fluor 488 in 2 % BSA for 2 h at room temperature. After hybridization, nuclei were counterstained using 1 μ g/mL DAPI for 10 min, and coverslips were mounted in Mowiol–based medium (10 % w/v Mowiol 4–88, 2.5 % DABCO, 25 % glycerol, 0.1 M Tris–Cl pH 8.5) and sealed with transparent nail polish.

##### b) Combined RNA FISH and immunofluorescence

For experiments involving both RNA FISH and antibody staining, coverslips were rinsed three times in 4x SSC following hybridization, blocked in 2% BSA/PBS for 2 h at room temperature, and processed for immunofluorescence as described previously (Maharana et al., 2023).

#### Image processing and quantification

Representative images were assembled in Fiji, applying uniform linear adjustments to brightness and contrast. To visualize stress–granule colocalization against intense mitochondrial RNA fluorescence, ∼10 % of pixels in the ND5 channel were intentionally saturated. Final panels were saved as JPEGs for presentation.

### Puromycylation assay for measuring translation rates

To assess translation activity under different stress conditions, ∼0.04 × 10^6^ HeLa cells were plated onto 12 mm No. 1 glass coverslips in 24–well plates and cultured overnight. During the final 10 min of each stress treatment cells were incubated with 10 μg/mL puromycin in complete medium. Cells were then fixed in 4 % formaldehyde for 10 min and permeabilized with 0.25 % Triton X–100 in 1x PBS for 10 min at room temperature. The puromycin incorporated in the nascent polypeptide chains was detected by immunostaining with anti–puromycin primary antibody (1:250 in 2 % BSA/1x PBS), followed by Alexa Fluor 488–conjugated secondary antibody (1:1000 in 2 % BSA/1x PBS) for 2 h in the dark at room temperature. Nuclei were counterstained with 1 μg/mL DAPI, and coverslips were mounted as described in the immunostaining protocol. Imaging was performed using a 60x/1.518 NA oil–immersion objective on a Nikon ECLIPSE Ti2 microscope.

### Fixed–cell imaging

Most fixed–cell images were acquired on a Nikon ECLIPSE Ti2 microscope equipped with a 60x/1.518 NA Type–F oil–immersion objective. Illumination was provided by a CoolLED PE–800 light source using the following excitation wavelengths: 405 nm (DAPI), 488 nm (GFP), 561 nm (mCherry), and 647 nm (far–red). Emission was collected using bandpass filters centered at 470 nm (GFP), 550 nm (mCherry), and 635 nm (far–red). Images were captured with an ORCA–Fusion BT camera (2304 × 2304 effective pixels; 6.5 μm pixel size). Confocal imaging of dsRNA and mitoGFP was performed on a Dragonfly 400 spinning–disk confocal system (Andor), equipped DFLY–CAM1–S2BV11. SONA 2.0B–11 camera. The images were acquired using a multi–line laser module (LM–405–100, 405 nm laser; LM–488–50, 488 nm laser; LM–561–50, 551 nm laser; and LM–637–140, 488 nm laser) on a 63x/1.40–0.60 oil objective. we also used sample RO spinning disk mounted on Nikon ECLIPSE Ti2 microscope with ORCA–Fusion BT camera. Instrument parameters—including laser power, exposure duration, and detector gain—were kept identical across all samples within each experiment to ensure consistent image acquisition.

### Super-resolution microscopy with DNA-PAINT

#### Imagers Fluorophore Conjugation and Purification

Imager DNA strands were conjugated with Cy3B as described earlier (*14*). In brief, imager sequences (Table 3) carrying an amine modification at the 3^’^-end were purchased from Merck and dissolved in Milli-Q® water at a concentration of 1 mM before being stored at -20 ^∘^C until conjugation. Fluorophore Cy3B-MonoNHS-Ester was dissolved in DMSO at a concentration of 13 mM and stored at -20 ^∘^C until conjugation. A 15 nanomoles of DNA was used during every conjugation reaction in a buffer containing 0.1 M NaHCO3 in 1× PBS. Five molar excess of fluorophore was added, vortexed vigorously, and left on a shaker at 4 ^∘^C overnight. The labelled imager was then purified using a reverse-phase column on an HPLC (Agilent Technologies). The purified product was then lyophilized and dissolved in Milli-Q® water before being stored at -20 °C for future use.

**Table 2:**
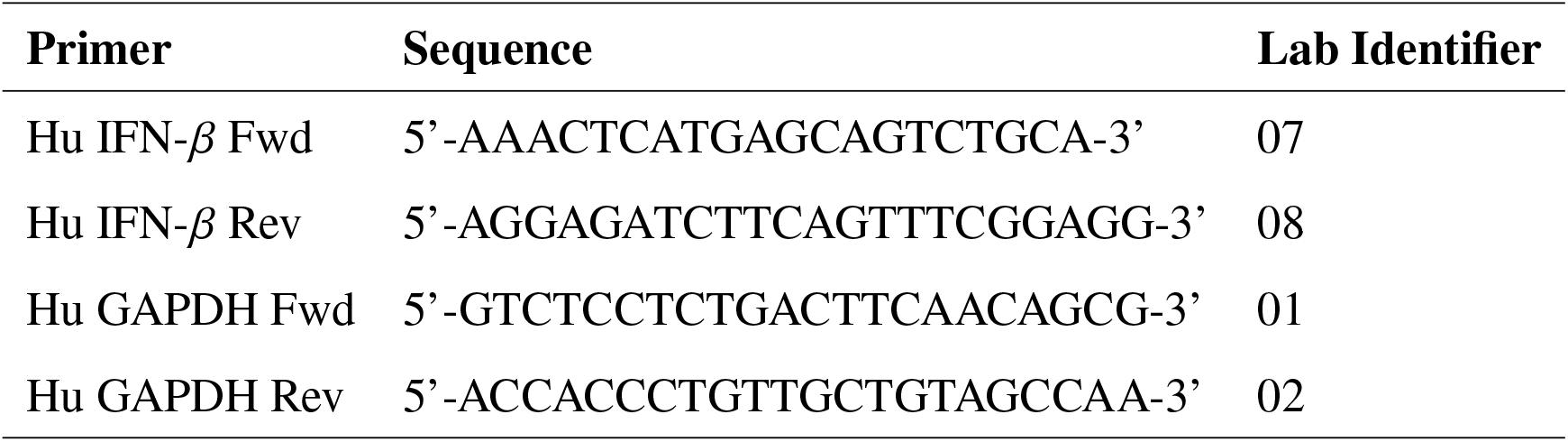
qPCR primers used in this study.

**Table 3:**
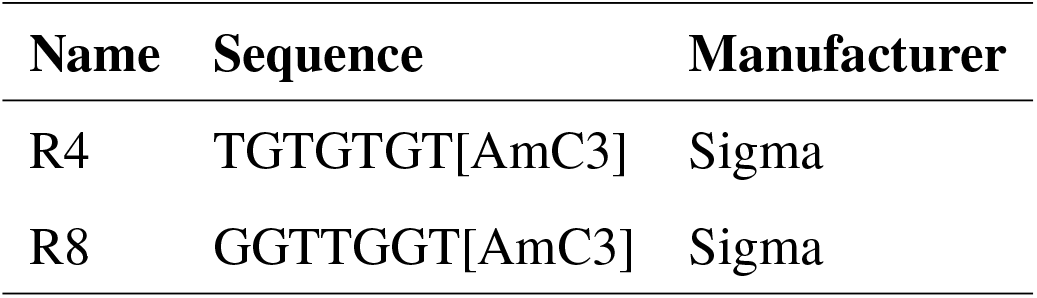
Imager Sequences.

#### Antibody – DNA Conjugation

Secondary antibody conjugation was performed as mentioned earlier (*15*). In brief, antibodies against rabbit IgG and mouse IgG were buffer exchanged to 1× PBS and concentrated to over 1.5 mg/ml using centrifugal concentrators. Hetero-bifunctional crosslinker DBCO-PEG4-NHS Ester was dissolved in DMSO and stored at a concentration of 15.39 mM at -80 ^∘^C. A 25-fold excess of crosslinker was added to 100 µg of antibody, and the mixture was incubated at ambient temperature for 2 hours with mild shaking. Excess crosslinker was removed by buffer exchange using a centrifugal filter till the crosslinker was diluted by 1000-fold. Conjugation ratio was measured using a NanoPhotometer® (Implen NP80) with absorbance measured at 260 nm, 280 nm, and 309 nm. A conjugation ratio of 9 cross-linkers to every antibody was achieved. Docking strands containing a 5’-C3-azide moiety (Table 4) were purchased from Metabion and dissolved in MilliQ® water at a concentration of 10 mM before being stored at -20 ^∘^C prior to use. A 15× excess of the docking strand was added to the cross-linker-containing antibody and incubated overnight on a shaker at 4 ^∘^C. Excess docking strands were removed using centrifugal filter, and the DNA-conjugated antibodies were stored in 4 ^∘^C until further use.

**Table 4:**
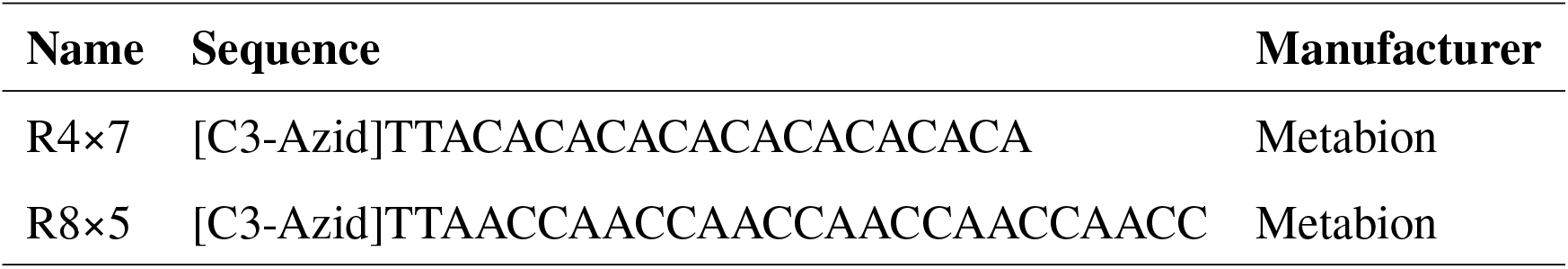
Docking strand Sequences.

#### Cell fixation and staining

Cells were cultured in cover-glass bottom dishes at a density of 8000 cells per well. Cells were then treated for oxidative stress or translation inhibition for the mentioned time. Prior to fixation, PFA fixation buffer (4 % paraformaldehyde in 1× PBS) was prewarmed to 37 °C on a heat block. The media was removed, and the prewarmed fixative was added immediately to the wells. The dish was moved to an incubator maintained at 37 °C for 10 minutes to minimize thermal fluctuations experienced by cells. Cells were then rinsed 3 times with 1× PBS, followed by incubation in blocking buffer (3% w/v BSA, 0.05 mg/ml Salmon Sperm DNA and 0.25% v/v Triton X-100) for 1 hour. Cells were then rinsed 3 times with 1× PBS prior to incubation with primary antibodies (G3BP1:#ab56574, 1:500 dilution; Tom20: Santa Cruz #SC-17764, 1:500 dilution) in antibody incubation buffer (3 % w/v BSA, 0.05 mg/ml Salmon Sperm DNA, 0.02 % v/v Tween-20 and 1 mM EDTA in 1× PBS) for 1 hour. Following this, cells were rinsed thrice with 1× PBS for 5 minutes each. Secondary antibodies conjugated to DNA strands were incubated in antibody incubation buffer at a 1:200 dilution for 45 minutes. Cells were rinsed thrice with 1× PBS for 5 minutes each. Cells were incubated with gold nanoparticles at a 1:10 dilution in buffer C (1× PBS supplemented with 500 mM NaCl) for 10 minutes, then washed 3 times with 1× PBS. These nanoparticles act as fiducial markers for drift correction and channel alignment.

#### Microscopy and Imaging

Imaging was performed on a Nikon Ti2 Eclipse microscope, mounted with a motorised H-TIRF unit and a perfect focus system (PFS). A Teledyne Photometrics PRIME BSI sCMOS camera was used to capture the images. The L6cc laser combiner from Oxxius Inc., France, provided illumination with a 561 nm. An oil-immersion high-NA objective lens capable of Total Internal Reflection Fluorescence (TIRF) imaging from Nikon (Nikon #Apo SR HP TIRF 100×, 1.49 NA, oil immersion) was used. The camera settings included 2×2 pixel binning and cropping to achieve a 66.56 × 66.56 µm^2^ field of view with each pixel measuring 130 nm × 130 nm. A 20 × PCD (Protocatechuate 3,4-dioxygenase) was prepared in 50 mM KCl, 1 mM EDTA, and 100 mM Tris-Cl, pH 8.0, and 50 % glycerol to a concentration of 6 µM. The stock PCD was then divided into 10 μl aliquots and stored at -20 °C for future use. A 40 × PCA (Protocatechuic Acid / 3,4-Dihydroxybenzoic acid) was made by dissolving 154 mg of PCA in 8 ml of Milli-Q®, and 3 M NaOH was added dropwise and stirred until PCD completely dissolved. Volume was made up to 10 ml, and 10 μl aliquots were stored at -20 ^∘^C for future use. A 100× Trolox (6-hydroxy-2,5,7,8-tetramethylchroman-2-carboxylic acid) was prepared by dissolving 100 mg of Trolox in 430 μl of methanol, 345 μl of 1 M NaOH, and 3.2 ml of Milli-Q®. The solution was stored as 20 μl aliquots at -20 °C for future use. 200 μl of imaging buffer was prepared in buffer C containing 1× PCA, 1× PCD, and 1× Trolox, along with the imager. Exchange-PAINT modality was used for two-channel imaging (*16*). Imaging parameters are outlined in Table 5. Image reconstruction and visualization Images were acquired using Nikon’s NIS Elements software. Images were stored as *.nd2 files and were reconstructed using the Localize module on Picasso (*17*). Drift correction and channel alignment were performed using redundant cross-correlation (RCC) followed by picked fiducial markers on the Render module in Picasso.

**Table 5:** Imaging parameters.

| Target | Prim. Ab | Sec. Ab | Img. | Conc. | Laser<br>(W/cm<sup>2</sup>) | Frames | Exp.<br>(ms) |
| --- | --- | --- | --- | --- | --- | --- | --- |
| Tom20 | SC-17764 | 715-005-150 | R8 | 250 pM | 250 | 10000 | 100 |
| G3BP1 | 61559 | 711-005-152 | R4 | 500 pM | 250 | 10000 | 100 |

### Image Analysis and Representation

#### Fixed–cell image analysis

Quantification of mean fluorescence intensities (for markers including G3BP1, G3BP2, poly(A)– tailed mRNAα, p–EIF2α, dsRNA, TOM20, p–PKR/PKR, DRP1, p–DRP1, and p–IRF3) was performed using a custom MATLAB script (R2024b, 24.2.0). Cellular compartments used for measurement are indicated above the corresponding graphs in the figures. Segmentation of specific cellular compartments was carried out in MATLAB, and each treatment condition was normalized to its respective vehicle control. Normalized values were visualized as scatter plots showing mean ± SD using RStudio (2024.12.0+467). Representative images were processed in Fiji, where brightness and contrast were adjusted linearly and uniformly across all samples. For intensity–based visualization, the Fire lookup table (LUT; low intensity = black, high intensity = yellow/white) was applied to emphasize the distribution of pixel intensities. Final images were exported as JPEG files for presentation.

#### Live–cell analysis of stress granule properties

Best–focus frames from time–lapse acquisitions were compiled into multi–page TIFF stacks. Drift correction was performed using the “Linear Stack Alignment with SIFT” plugin in Fiji. The drift–corrected stacks were then analyzed using MATLAB (R2024b, 24.2.0). To characterize SG dynamics, parameters including SG area fraction, mean fluorescence intensity of mCherry–G3BP1 within SGs, and the number of SGs per cell were extracted using custom MATLAB pipelines. Automated segmentation was applied to identify SG structures; mCherry–G3BP1 puncta smaller than 0.0049 ^2^ (corresponding to ∼250 nm diameter) were excluded from analysis. SG–positive cells were counted manually in Fiji using the Cell Counter plugin (*13*). For SG area fraction and SG number per cell, raw values were plotted directly. G3BP1 enrichment (MFI_SG_/MFI_cell_) was normalized using min–max normalization:

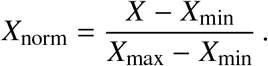

Plots were generated in RStudio representing mean ± SD. Representative images were prepared in Fiji with linear adjustments to brightness and contrast across all conditions and time points. The ICA LUT was used for intensity–based displays. Final panels were saved as JPEGs for presentation.

### Optogenetic induction of stress granules and inflammatory signaling

HeLa cells were subjected to optogenetic SG induction as described above. Thirty minutes after activation, cells were fixed in 4% formaldehyde and immunostained for phospho–IRF3 (p–IRF3) following the immunofluorescence protocol. Nuclear and cytoplasmic compartments were segmented to compute p–IRF3 enrichment as the ratio of nuclear mean fluorescence intensity (MFI) to cytoplasmic MFI using a custom–written MATLAB script. In the same cells, mCherry–G3BP1 mean fluorescent intensity was also quantified. **i) Spectral leakage correction for representative images**

To correct bleed–through from the mCherry channel into far–red, we estimated the channel leakage fraction (*λ*) using mCherry–only controls acquired with far–red settings. Far–red images were corrected in Fiji according to:

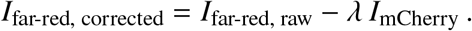

Brightness and contrast were adjusted linearly and uniformly across conditions for presentation. **ii) Regression and ANCOVA**

The relationship between optoS/G3BP1 burden and p–IRF3 enrichment was assessed by ordinary least–squares linear regression, with p–IRF3 enrichment as the dependent variable and normalized mCherry–G3BP1 intensity as the predictor using a custom–written MATLAB code. Scatter plots display individual cells, fitted regression lines per group, 95 % confidence bands, and in–panel reporting of the regression equation (Fig. 4C). Group comparisons (G3BP1 vs opto–G3BP1) were evaluated by analysis of covariance (ANCOVA). The significant difference between the two groups was reported as the slope–difference p value.

## Mitochondrial morphology analysis

### Sample preparation for fixed cell imaging

Approximately 0.3 million HeLa cells were seeded onto 12 mm. 1 thickness glass coverslips and cultured overnight at 37 °C under 5 % CO_2_ in a humidified incubator. After 12 hours, the cells were transfected with the Mito–GFP plasmid using X–Fect transfection reagent according to the manufacturer’s protocol. The cells were further allowed to recover in high–glucose DMEM media containing 5 % FBS for 12–16 hours. Coverslips containing live cells were treated with the desired drug or vehicle control and fixed using 4 % formaldehyde for 10 min at 37 ∘C. The coverslips were then permeabilised with 0.25 % Triton X–100 in 1x PBS. Following permeabilisation, coverslips were stained with 1 μg/mL DAPI for 10 minutes at room temperature in a humidified chamber and mounted onto microscope slides using mounting media.

### Image acquisition for fixed cell imaging and 3D mitochondrial morphology analysis

The mounted coverslips were imaged using a 63x/1.40 NA oil–immersion objective on a Leica microscope, connected to a Dragonfly 400 spin–disk confocal microscope (Andor). A multi–line laser module (LM–488–50, 488 nm laser) was used as a source of excitation wavelength 488 nm (mitoGFP), and emission was collected using 445/46 325 mm diameter bandpass. DFLY–CAM1– S2BV11. SONA 2.0B–11 camera with a resolution of 2304 × 2304 effective pixels and a pixel size of 6.5 μm x 6.5 μm was used to acquire images. To capture the mitochondria 3D network, images were acquired in a z–stack of 17–20 frames with a 0.4 μm step size covering the whole cell.

### Sample preparation for live cell imaging and image acquisition for 2D mitochondrial morphology analysis

HeLa ωDRP1 cells being thin in the z–resolution, favored live–cell imaging over time for mitochondrial morphology analysis. About 0.3 million cells were seeded on 22 mm No. 1 thickness glass coverslips and allowed to grow overnight at 37 °C and 5 % CO_2_ in a humidified incubator. The following day, cells were transfected with the mitoGFP plasmid as previously described. The next day, coverslips were mounted on steel live cell dishes in cell culture media containing 10 mM HEPES. The imaging was performed on live cells incubated at 25 °C on the temperature– controlled microscopy incubator of the Nikon ECLIPSE Ti2 microscope using a 60×1.518 Type F oil–immersion objective. The desired treatment was induced on the microscopy stage, and the cells were imaged over time using a 488 nm excitation wavelength in the best focus plane.

### Mitochondrial morphology analysis

Mitochondrial morphology of the fixed samples was analysed in 3D using the 3D analysis option in Mitochondrial Analyzer plugin in ImageJ/FIJI, available at https://github.com/AhsenChaudhry/Mitochondria-Analyzer. The analysis required thresholding the images for proper mitochondrial masking in 3D and 2D.

#### 2D/3D Image Thresholding

Images were thresholded using the local thresholding method in the mitochondria analyser plugin in FIJI. In the local threshold method, Weighted Mean was selected for all image sets. For each experimental condition and cell, the block size, C–value, and outline radius were optimised based on visual assessment of the original and thresholded images. The rest of the default settings were kept unchanged. To verify the accuracy of mitochondria thresholding in different mitochondrial morphology states, we transfected mitochondria–localized GFP or mitoGFP in HeLa cells with DRP1 KO (HeLa ωDRP1) to generate elongated mitochondria (*18*), or treated cells with CCCP, known to induce mitochondrial fragmentation (*19*). This was compared with untreated HeLa WT cells, which have a mixed morphology (Fig. S7A).

#### 2D/3D Image analysis

Mitochondrial morphology was quantified using the 3D and 2D analysis options in the Mitochondrial Analyzer. For 3D mitochondria morphology analysis, mean branch length, sphericity, mean surface area, and mean volume were calculated per mitochondrion per cell across all z planes (Fig. S7B). The elongated mitochondria showed a 40% increase in mean branch length, whereas CCCP treatment resulted in a 23.7% decrease (Fig. S7C). The sphericity of mitochondria increased as they fragmented, from 0.09 in elongated mitochondria to 0.11 in intermediate mitochondria and finally to 0.37 in fragmented mitochondria (Fig. S7D). Similar to mean branch length, mean surface area, and mean volume, these parameters were also higher in elongated mitochondria and decreased from intermediate length to fragmented mitochondria as the morphology became more fragmented (Fig. S7, E and F). For live–cell 2D analysis of HeLa ωDRP1 cells at the best focus plane, single cells were tracked over time. Parameters including mean branch length, aspect ratio, mean perimeter, and mean area were calculated at each time point per cell. The difference between t = 0 and t = n (final time point) was plotted per cell to show parameter differences at t = n relative to t = 0 (Fig. S10, B–E).

### RNA Isolation and Quantitative RT–qPCR

Total cellular RNA was extracted using TRIzol. Cells were lysed directly in TRIzol reagent, after which 0.2 mL chloroform was added per 1 mL TRIzol. Samples were mixed thoroughly and centrifuged at 12,000 g for 15 min at 4 °C. The aqueous phase was transferred to a fresh tube, and RNA was precipitated by adding 0.5–1 mL of isopropanol and incubating the mixture at room temperature for 10 min. Following centrifugation at 12,000 g for 10 min at 4 °C, the RNA pellet was washed with 75 % ethanol and centrifuged at 7,500 g for 5 min at 4 °C. After air–drying the pellet for ∼10 min, RNA was resuspended in 25 μL RNase–free water supplemented with 200 U/mL RNase inhibitor (RRI). RNA concentration and purity were assessed using a NanoDrop– 1000 spectrophotometer. cDNA synthesis was performed using the PrimeScript™ cDNA Synthesis Kit according to the manufacturer’s instructions. Quantitative RT–qPCR was carried out using TB Green Premix Ex Taq™ II on a QuantStudio 6 Real–Time PCR System (Applied Biosystems) following the supplier’s protocol. Gene expression levels were normalized to GAPDH, and relative transcript abundance was calculated using the ωωCt method. Primer sequences used in this study are listed in Table 2.

### Polysome profiling

HeLa cells cultured in 100 mm dishes were first treated with cycloheximide (100 μg/mL) for 10 min at 37 °C followed by individual inhibitor treatment: puromycin (7 μg/mL), and emetine (40 μM) for 30 min. The cells were processed further on ice. Cells were first washed with ice–cold 1x PBS containing cycloheximide and then with ice–cold hypotonic buffer [5 mM Tris–HCl (pH 7.5), 1.5 mM KCl, 5 mM MgCl_2_, and 100 μg/mL cycloheximide]. Cells were scraped in ice–cold lysis buffer [5 mM Tris–HCl (pH 7.5), 1.5 mM KCl, 5 mM MgCl_2_, 100 μ g/mL cycloheximide, 1 mM DTT, 200 U/mL RNasin, 200 μg/mL tRNA, 0.5 % Triton X–100, 0.5 % sodium deoxycholate, and 1x protease inhibitor cocktail], vortexed briefly and incubated on ice for 15 min. The lysate KCl concentration was adjusted to a final concentration of 150 mM, followed by centrifugation at 3,000 x g for 10 min at 4 °C. The resulting supernatant was collected and used immediately or flash frozen and stored at −80 °C for later use. Subsequently, 3 OD_254_ of total RNA was loaded on top of a 10 %–50 % sucrose gradient and ultracentrifuged at 36,000 rpm for 2 h at 4 °C using an SW41 rotor (Beckman). Finally, polysome profiles were visualized using a polysome profiler (BioComp, model number). The polysome profiles were plotted using RStudio (2024.12.0+467).

### Western blotting and analysis

#### Protein isolation and quantification

About 0.5 million cells were cultured in a 6–well cell culture plate overnight. The following day, cells were trypsinized, and cell pellets were lysed using 100 μL of lysis buffer containing 20 mM Tris–HCl (pH 7.5), 1 mM Na_2_EDTA, 150 mM NaCl, 1 mM EGTA, 1 % Triton X–100, 2.5 mM sodium pyrophosphate, 1 mM Na_3_VO_4_,1 μg/mL leupeptin, and 1 mM *β*–glycerophosphate. Lysates were vortexed for 30 s, then incubated on ice for 5 min. This step was repeated 6 times to effciently lyse the cells. Lysates were centrifuged at 13,000 rpm for 10 min at 4 °C, and supernatants were collected. Bradford assay was performed to determine total protein concentration in the lysates. Briefly, a serial dilution of BSA solution (1 mg/mL) ranging from 0.05 to 0.5 mg/mL was first assayed to generate a standard curve of protein concentration vs. absorbance at 595 nm. The concentration of the unknown protein sample was determined by adding 200 μL of a 1:5 dilution of Bradford reagent and Milli–Q water. After 10 min of incubation in the dark, the protein absorbance of each sample was measured at 595 nm using a Tecan plate reader. Protein concentration was derived from the absorbance value using the BSA standard curve.

#### SDS–PAGE and immunoblotting

For SDS–PAGE, equal amounts of protein were mixed with 1x loading dye containing 250 mM Tris–Cl (pH 6.8), 8% SDS, 0.1% bromophenol blue, 40% v/v glycerol, and 100 mM DTT and denatured at 95 °C for 10 min. Protein samples were resolved in 8% SDS–PAGE gel. Proteins were transferred to a nitrocellulose membrane using wet transfer (buffer containing 25 mM Tris base, 20 % methanol, and 190 mM glycine) for 2–3 h at room temperature. The membrane was blocked with 5 % BSA in 1x TBST buffer for 1 h at room temperature, then incubated with primary antibody dissolved in 5 % BSA/1x TBST overnight at 4 °C. The following day, the membrane was washed times with 1x TBST and incubated with an HRP–conjugated secondary antibody for 1 h at room temperature. For signal detection, the membrane was incubated with Clarity™ ECL substrate for a few seconds, followed by chemiluminescence detection using a Bio–Rad Chemidoc imager.

#### Data analysis and plotting

FIJI was used to calculate protein band intensities. Each band was normalised to GAPDH within the same sample. Data were plotted as bar graphs using RStudio (2024.12.0+467).

### Subcellular fractionation and ERMCS isolation

ERMCS were isolated as previously described (*20*). Briefly, 2.5 million HeLa cells were seeded into a 100 mm plate and incubated for 48 hours. After this, 15 such plates per condition, at around 80 % confluency, were trypsinized, pelleted, and washed with 1x PBS. All subsequent steps were performed on ice at 4 °C in buffers containing protease inhibitor cocktail. The cell pellet was lysed using IB_cells_-1 lysis buffer (30 mM Tris–HCl, 225 mM mannitol, 75 mM sucrose, and 0.1 mM EGTA, pH 7.4) for a few minutes on ice, then centrifuged at 700 g to remove cell debris and the nuclear fraction. The supernatant (post–nuclear lysate) was collected and further centrifuged at 7000 g twice. The pellet from this spin contained mitochondria, whereas the supernatant consisted of plasma membrane, lysosomes, ER, and cytosol. The mitochondrial pellet was resuspended in IB_cells_-2 (30 mM Tris–HCl, 225 mM mannitol, 75 mM sucrose) and further centrifuged at 10,000 g to remove organellar contamination. The crude mitochondrial pellet was then resuspended in MRB buffer (250 mM mannitol, 5mM HEPES, 0.5mM EGTA-pH7.4)and subjected to ultracentrifugation (Beckman Coulter Optima L–90K Ultracentrifuge). To separate pure mitochondria and ERMCS, the crude mitochondria were loaded onto a Percoll gradient (30 % Percoll [v/v]) containing 225 mM mannitol and 25 mM HEPES (pH 7.4) in the ultracentrifuge tube (Beckman Coulter, #344060) and centrifuged at 95,000 g (27,500 rpm with a SW40–Ti Beckman Coulter rotor) for 30 min at 4 °C. The ERMCS layer was collected and centrifuged at 6300 g for 10 min. ERMCS were further pelleted by centrifuging the supernatant collected in the previous step at 100,000 g (28,200 rpm with a SW40–Ti Beckman Coulter rotor). The pellet containing ERMCS was resuspended in RIPA buffer (50 mM Tris–HCl; pH 8.0, 150 mM NaCl, 1 % NP40, 0.1 % SDS, 0.5 % sodium deoxycholate) and processed for western blotting as mentioned previously.

### Fluorescence correlation spectroscopy (FCS)

FCS was done on HeLa WT or HeLa ωDRP1 cells seeded on a 22 mm glass coverslip dish and transiently transfected with mCherry–G3BP1 using PEI. For FCS, the cells were maintained in complete cell culture media containing 10 mM HEPES. Confocal FCS was performed on a SteadyCam optical setup (Abberior Instruments GmbH) integrated with a Leica SP5 microscope at room temperature. For confocal FCS, a pulsed 561 nm Laser (output power 0.2 mW, repetition rate 40 MHz, pulse length *<* 100 ps, pulse energy *>* 3 pJ) using a 100x/1.4 NA STED WHITE objective was used to excite the mCherry–G3BP1 protein, and the emission was recorded in 575–625 nm using a Picoharp 300 TCSPC module (PicoQuant) and SymPho Time 64 software (PicoQuant), and LSM upgrade toolkit (Leica). Cells expressing mCherry–G3BP1 were selected, and the laser was parked in the cytoplasm showing a diffused mCherry–G3BP1 signal (Fig. 4C, white inset). FCS readings were collected at time 0, after which Puro or SA was added. FCS readings from 15–20 transfected cells across 3 biological replicates were collected for up to 45 min after treatment. The FCS data were binned for 10–min intervals, with bin centres at 5, 15, 25 and 35 min for further analysis. For a single FCS reading, the laser was focused on the confocal volume within the cell cytoplasm for 10–15 s to measure intensity fluctuations arising from fluorescent molecules entering and leaving the illumination area. Repeats were performed across the entire FCS measurement to assess reproducibility. FCS analysis was performed in QuickFit software (Langowski Group (B040), DKFZ, http://www.dkfz.de/Macromol/quickfit/) using a 3D anomalous diffusion model (*21*). The autocorrelation function (Fig. 4D) was calculated from the intensity time trace and fitted using the equation below (*22, 23*):

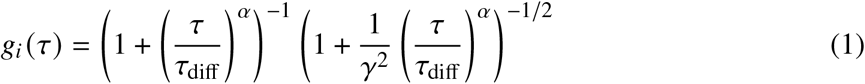

where α is the anomality parameter, τ_diff_ is the diffusion time and *γ* is the aspect ratio = 6.

From the fitted G(τ), the transit time was calculated. We calculated the diffusion constant D from the transit time using the appropriate model (*24*).

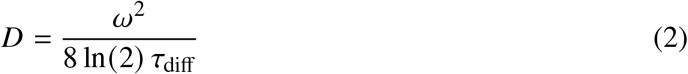

Where, ω is the FWHM of the confocal volume in x-y plane = 285 nm. To calculate particle size from *D*, we measured the cytoplasm viscosity (see section below).

### Viscosity calibration, cell viscosity measurement and size estimation

To calculate the size of the diffusing species (protein hydrodynamic radius) using the Stokes– Einstein equation, we require viscosity of the cell cytoplasm. We measured the microscopic viscosity of the cell cytoplasm using a fluorescent molecular rotor, Bodipy C–12, whose lifetime depends on the viscosity of the surrounding medium. The relation between lifetime (τ) and viscosity (η) can be calculated using the Förster–Hoffmann relation (*25, 26*):

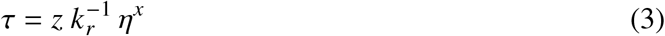

Where z and x are constants and *k*_*r*_ is the radiative rate constant. From the viscosity calibration, we obtained z/ *k*_*r*_ = 0.455 and x = 0.324.

The lifetime needs to be calibrated with solutions of known viscosities. We used a series of glycerol and methanol mixtures at various concentrations (from G/M 0/100 to G/M 100/0) with 0.5% Bodipy C–12 (*27*). These solutions were imaged through a cover glass–bottomed sample chamber. Fluorescence lifetime data acquisition was performed using SymPhoTime 64 software, and fluorescence lifetimes were determined through Time–Correlated Single Photon Counting (TCSPC) using a Stedycon optical setup (Abberior Instruments GmbH) integrated with a Leica SP5 microscope. In TCSPC, the time between excitation of the sample by the 488 nm pulsed laser and arrival of the emitted photon at the detector was measured. To generate a fluorescence lifetime image, photons were attributed to distinct pixels by recording both the absolute photon arrival time and the relative arrival time with respect to the laser pulse. FLIM images were captured with a frame size of 30 × 30 μm.

FLIM images were analyzed using several steps to ensure accurate data interpretation. First, a binning factor of 2 was applied to combine adjacent pixels into larger super–pixels to improve signal–to–noise ratio without significantly compromising spatial resolution. We utilized the “n– exponential tail fit” model (*28*) to fit the FLIM data (Supp. Fig. 4.1A).

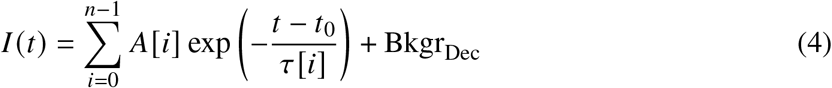

The known viscosities of glycerol and methanol were plotted against the fitted average life-time values. Data were fitted using the Förster–Hoffmann equation to obtain calibration parameters (Supp. Fig. 4.1B and 4.1C). FLIM measurements were then performed on Bodipy C–

12 stained HeLa WT cells and analyzed similarly to map cytoplasmic lifetime into viscosity (Supp. Fig. 4.1D–F). Bright regions were intentionally ignored to remove inherent dye clusters. Custom ROIs were used to select cytoplasm for final FLIM analysis. Fits and residuals are shown in Supp. Fig. 4.1G and 4.1H. Lifetime values and corresponding viscosity values were plotted as histograms (Supp. Fig. 4.1I–J). Inside cell the viscosity have a mean value around 94.79 and SD 51.17 Pa.S (57% of the mean).

After obtaining viscosity values and the previously calculated *D* values, the mean diameter (d) of the diffusing species was calculated using the Stokes–Einstein equation (*29, 30*):

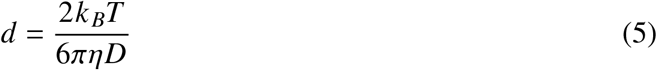

### Error analysis in calculating diameter of the nanoSGs

The error in calculating diameter (d) of the nanoSGs comes due to two independent measurements of diffusivity (D) and viscosity (*η*). The heterogeneous distribution of the D and *η* contributes to the error in these quantities. Assuming the measurements for viscosity (*η*) and diffusion (*D*) are independent and random, their relative errors add in quadrature (the square root of the sum of the squares).

The relative error in the diameter (*δd*) is calculated as:

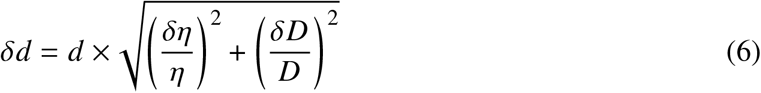

where *δ* _*η*_ and *δ*_*D*_ are the SD of *η* and D. This computed error is added to the computed nanoSG sizes shown in figure Fig. 4G.

Statistical comparisons between independent groups were performed using the non-parametric, two-sided Mann-Whitney U test as discussed previously. As the viscosity measurements were performed independently, the estimation of nanoSG diameter from the diffusivity measurements also contains the error in viscosity, due to viscosity variation in cell. Now, the relative error in individual value of d only due to viscosity will be Δ*d* = *d* × (Δ*η*/ *η*). So, to robustly account for the individual measurement uncertainty (±57% as shown in the previous section) associated with each data point of d, a Monte Carlo simulation approach was implemented.

For each iteration of the Monte Carlo simulation, the value of every individual data point, d, was randomly sampled from a uniform distribution bounded by its specific error margins [*d* − 0.57 *d, d* + 0.57 *d*]. The Mann-Whitney U test was subsequently performed on these randomized arrays. This resampling and testing process was repeated for 10,000 iterations per group comparison to generate a distribution of simulated *p*-values.

To establish the absolute limits of statistical variance within the error bounds, the “best-case” and “worst-case” *p* -values were extracted as the lower (2.5th percentile) and upper (97.5th percentile) bounds of the 95% confidence interval, respectively. In Fig. 4G, the “worst-case,” i.e., upper (97.5th percentile) bounds of the 95 % confidence interval are mentioned. Analyses are performed using custom Python scripts utilizing the *SciPy* and *NumPy* libraries.

## References and Notes

1. Y. Wu, et al., Translation landscape of stress granules. Science Advances 11 (40), eady6859 (2025).

2. S. Markmiller, et al., Context-dependent and disease-specific diversity in protein interactions within stress granules. Cell 172 (3), 590–604 (2018).

3. A. Khong, et al., The stress granule transcriptome reveals principles of mRNA accumulation in stress granules. Molecular cell 68 (4), 808–820 (2017).

4. S. Chen, J. Zhang, F. Zhao, Screening linear and circular RNA transcripts from stress granules. Genomics, proteomics & bioinformatics 21 (4), 886–893 (2023).

5. C. Bussi, et al., Stress granules plug and stabilize damaged endolysosomal membranes. Nature 623 (7989), 1062–1069 (2023).

6. S. Liu, et al., Mammalian IRE1α dynamically and functionally coalesces with stress granules. Nature cell biology 26 (6), 917–931 (2024).

7. Y. Zhang, J. Seemann, RNA scaffolds the Golgi ribbon by forming condensates with GM130. Nature cell biology 26 (7), 1139–1153 (2024).

8. T. Amen, D. Kaganovich, Stress granules inhibit fatty acid oxidation by modulating mitochondrial permeability. Cell reports 35 (11) (2021).

9. X. Qi, M.-H. Disatnik, N. Shen, R. A. Sobel, D. Mochly-Rosen, Aberrant mitochondrial fission in neurons induced by protein kinase Cδ under oxidative stress conditions in vivo. Molecular biology of the cell 22 (2), 256–265 (2011).

10. K. Momma, T. Homma, R. Isaka, S. Sudevan, A. Higashitani, Heat-induced calcium leakage causes mitochondrial damage in Caenorhabditis elegans body-wall muscles. Genetics 206 (4), 1985–1994 (2017).

11. P. Jain, Z.-Q. Luo, S. R. Blanke, Helicobacter pylori vacuolating cytotoxin A (VacA) engages the mitochondrial fission machinery to induce host cell death. Proceedings of the National Academy of Sciences 108 (38), 16032–16037 (2011).

12. C. Teodorof-Diedrich, S. A. Spector, Human immunodeficiency virus type 1 gp120 and Tat induce mitochondrial fragmentation and incomplete mitophagy in human neurons. Journal of virology 92 (22), 10–1128 (2018).

13. W. Song, et al., Mutant huntingtin binds the mitochondrial fission GTPase dynamin-related protein-1 and increases its enzymatic activity. Nature medicine 17 (3), 377–382 (2011).

14. J. Kim, et al., VDAC oligomers form mitochondrial pores to release mtDNA fragments and promote lupus-like disease. Science 366 (6472), 1531–1536 (2019).

15. H. Xian, et al., Oxidized DNA fragments exit mitochondria via mPTP-and VDAC-dependent channels to activate NLRP3 inflammasome and interferon signaling. Immunity 55 (8), 1370–1385 (2022).

16. A. Dhir, et al., Mitochondrial double-stranded RNA triggers antiviral signalling in humans. Nature 560 (7717), 238–242 (2018).

17. G. Kroemer, J. C. Reed, Mitochondrial control of cell death. Nature medicine 6 (5), 513–519 (2000).

18. S. Maharana, et al., SAMHD1 controls innate immunity by regulating condensation of immunogenic self RNA. Molecular cell 82 (19), 3712–3728 (2022).

19. M. Paget, et al., Stress granules are shock absorbers that prevent excessive innate immune responses to dsRNA. Molecular cell 83 (7), 1180–1196 (2023).

20. N. Curdy, et al., The proteome and transcriptome of stress granules and P bodies during human T lymphocyte activation. Cell reports 42 (3) (2023).

21. J. Smith, D. P. Bartel, The G3BP stress-granule proteins reinforce the integrated stress response translation programme. Nature Cell Biology pp. 1–14 (2025).

22. J. Guillén-Boixet, et al., RNA-induced conformational switching and clustering of G3BP drive stress granule assembly by condensation. Cell 181 (2), 346–361 (2020).

23. A. Wang, et al., Cancer cells evade stress-induced apoptosis by promoting HSP70-dependent clearance of stress granules. Cancers 14 (19), 4671 (2022).

24. P. Yang, et al., G3BP1 is a tunable switch that triggers phase separation to assemble stress granules. Cell 181 (2), 325–345 (2020).

25. O. Bounedjah, et al., Free mRNA in excess upon polysome dissociation is a scaffold for protein multimerization to form stress granules. Nucleic acids research 42 (13), 8678–8691 (2014).

26. Y. Kim, et al., PKR senses nuclear and mitochondrial signals by interacting with endogenous double-stranded RNAs. Molecular cell 71 (6), 1051–1063 (2018).

27. L. Zhang, et al., Mitochondrial double-stranded RNA drives aging-associated cognitive decline. Cell Research pp. 1–18 (2026).

28. V. López-Polo, et al., Release of mitochondrial dsRNA into the cytosol is a key driver of the inflammatory phenotype of senescent cells. Nature communications 15 (1), 7378 (2024).

29. S. Kim, et al., Mitochondrial double-stranded RNAs govern the stress response in chondrocytes to promote osteoarthritis development. Cell Reports 40 (6) (2022).

30. H. Lee, et al., Cell type-specific transcriptomics reveals that mutant huntingtin leads to mitochondrial RNA release and neuronal innate immune activation. Neuron 107 (5), 891–908 (2020).

31. E. McEwen, et al., Heme-regulated inhibitor kinase-mediated phosphorylation of eukaryotic translation initiation factor 2 inhibits translation, induces stress granule formation, and mediates survival upon arsenite exposure. Journal of Biological Chemistry 280 (17), 16925–16933 (2005).

32. T. Grousl, et al., Heat shock-induced accumulation of translation elongation and termination factors precedes assembly of stress granules in S. cerevisiae. PloS one 8 (2), e57083 (2013).

33. N. G. Farny, N. L. Kedersha, P. A. Silver, Metazoan stress granule assembly is mediated by P-eIF2α-dependent and-independent mechanisms. Rna 15 (10), 1814–1821 (2009).

34. S. C. Kamerkar, F. Kraus, A. J. Sharpe, T. J. Pucadyil, M. T. Ryan, Dynamin-related protein 1 has membrane constricting and severing abilities sufficient for mitochondrial and peroxisomal fission. Nature communications 9 (1), 5239 (2018).

35. S. Nagashima, et al., Golgi-derived PI (4) P-containing vesicles drive late steps of mitochondrial division. Science 367 (6484), 1366–1371 (2020).

36. J. A. Kashatus, et al., Erk2 phosphorylation of Drp1 promotes mitochondrial fission and MAPK-driven tumor growth. Molecular cell 57 (3), 537–551 (2015).

37. I. Zaja, et al., Cdk1, PKCδ and calcineurin-mediated Drp1 pathway contributes to mitochondrial fission-induced cardiomyocyte death. Biochemical and biophysical research communications 453 (4), 710–721 (2014).

38. Y. Wang, et al., Inhibition of PKR protects against H2O2-induced injury on neonatal cardiac myocytes by attenuating apoptosis and inflammation. Scientific reports 6 (1), 38753 (2016).

39. M. Dey, B. R. Mann, A. Anshu, M. A.-u. Mannan, Activation of protein kinase PKR requires dimerization-induced cis-phosphorylation within the activation loop. Journal of Biological Chemistry 289 (9), 5747–5757 (2014).

40. N. V. Jammi, L. R. Whitby, P. A. Beal, Small molecule inhibitors of the RNA-dependent protein kinase. Biochemical and biophysical research communications 308 (1), 50–57 (2003).

41. T. Cesaro, et al., PKR activity modulation by phosphomimetic mutations of serine residues located three aminoacids upstream of double-stranded RNA binding motifs. Scientific Reports 11 (1), 9188 (2021).

42. N. Kedersha, et al., G3BP–Caprin1–USP10 complexes mediate stress granule condensation and associate with 40S subunits. Journal of Cell Biology 212 (7), e201508028 (2016).

43. P. Zhang, et al., Chronic optogenetic induction of stress granules is cytotoxic and reveals the evolution of ALS-FTD pathology. elife 8, e39578 (2019).

44. N. Kedersha, et al., Dynamic shuttling of TIA-1 accompanies the recruitment of mRNA to mammalian stress granules. The Journal of cell biology 151 (6), 1257–1268 (2000).

45. Y. Adachi, et al., Drp1 tubulates the ER in a GTPase-independent manner. Molecular cell 80 (4), 621–632 (2020).

46. C. Duan, et al., Activated Drp1 initiates the formation of endoplasmic reticulum-mitochondrial contacts via shrm4-mediated actin bundling. Advanced Science 10 (36), 2304885 (2023).

47. N. More, J. Joseph, Disruption of ER–mitochondria contact sites induces autophagy-dependent loss of P-bodies through the Ca2+-CaMKK2-AMPK pathway. Journal of Cell Science 138 (5), JCS263652 (2025).

48. R. Zhou, A. S. Yazdi, P. Menu, J. Tschopp, A role for mitochondria in NLRP3 inflammasome activation. Nature 469 (7329), 221–225 (2011).

49. H.-R. Zhou, K. He, J. Landgraf, X. Pan, J. J. Pestka, Direct activation of ribosome-associated double-stranded RNA-dependent protein kinase (PKR) by deoxynivalenol, anisomycin and ricin: a new model for ribotoxic stress response induction. Toxins 6 (12), 3406–3425 (2014).

50. S. Frank, et al., The role of dynamin-related protein 1, a mediator of mitochondrial fission, in apoptosis. Developmental cell 1 (4), 515–525 (2001).

51. T. Duan, et al., Mito-tempo ameliorates tubular injury of diabetic nephropathy via inhibiting mt-dsRNA release and PKR/eIF2α pathway activation. Free Radical Biology and Medicine 237, 147–159 (2025).

52. A. Makino, B. Scott, W. Dillmann, Mitochondrial fragmentation and superoxide anion production in coronary endothelial cells from a mouse model of type 1 diabetes. Diabetologia 53 (8), 1783–1794 (2010).

53. T. Petrozziello, et al., Targeting tau mitigates mitochondrial fragmentation and oxidative stress in amyotrophic lateral sclerosis. Molecular neurobiology 59 (1), 683–702 (2022).

